# Chronobiological rhythms control of site- and cell-specific miRNA and mRNA genes and networks across the central nervous system

**DOI:** 10.1101/2024.09.09.612109

**Authors:** Amanda M. Zacharias, Ciara D. O’Connor, Danai G. Topouza, Zhi Yi Fang, Helia Ghazinejad, Hanlin Chen, Qingling Duan, Nader Ghasemlou

**Author notes:** These authors contributed equally (Qingling Duan, Nader Ghasemlou). **Corresponding authors:** *Nader Ghasemlou, PhD* 18 Stuart Street, room 754 Botterell Hall, Queen’s University Kingston, Ontario, Canada K7L 3N6 e *Qingling Duan, PhD* 18 Stuart Street, room 944 Botterell Hall, Queen’s University Kingston, Ontario, Canada K7L 3N6.

## Abstract

Biological rhythms control gene expression, but effects on central nervous system (CNS) cells and structures remain undefined. While circadian (24-hour) rhythms are most studied, many genes have periods of greater and less than 24-hours; these fluctuations can be both site- and cell-specific. Identifying patterns of gene rhythmicity across the CNS is necessary for both the study of chronobiology and to make sense of data obtained in the laboratory. We now identify cycling mRNAs, miRNAs, gene networks and novel mRNA-miRNA co-expression pairs in the cortex, hypothalamus, and corpus striatum using high-dimensional datasets. A searchable catalogue (https://www.ghasemloulab.ca/chronoCNS) was created to help refine the analysis of cellular/molecular rhythmicity across the CNS. Immunofluorescence was also used to confirm the rhythmicity of key targets across cells in these structures, with strong cycling signatures in resting oligodendrocytes. Our study sheds light on the contribution of circadian, ultradian, and infradian rhythms and mRNA-miRNA interactions to CNS function.

## Main

Circadian rhythms are 24-hour endogenous processes that help organisms adapt to the transition between day and night. Transcription-translation loops govern these processes through clock genes, which regulate the expression of genes in a tissue- and cell-specific manner to drive various outcomes^1^. Emerging evidence from naïve and disease states suggest that rhythms are integral to CNS function and activity. For example, the hypothalamus controls circadian activity via hormone secretion^2^; this rhythmicity affects outcomes including cognition and memory (e.g., in the cerebral cortex)^3^, and goal-directed behaviour (e.g., in the striatum)^4^. These sites not only have distinct functions, but also highly specific gene expression that represent over 5,000 cell clusters^5^. Gaining an understanding of when and where gene expression changes across the naïve CNS is fundamental to our understanding of neurobiological outcomes; therefore, there is a need for more detailed profiling of gene rhythmicity and gene-gene interactions across CNS sites.

Circadian (24-hour) rhythms govern most cycling genes, though ultradian (<24-hour) and infradian (>24-hour) periods are also possible^6^. Indeed, a foundational study estimated that ∼ 40% of the mouse protein-coding genome has a circadian rhythm in at least one of 12 organs – including the brainstem, cerebellum, and hypothalamus –though only 10 genes were determined to be cycling across all organs, of which seven were core clock factors^7^. These rhythms are governed by the central suprachiasmatic nucleus of the hypothalamus. Interestingly, it was found that 16% of genes in the liver (considered to be among the most rhythmic organs) were under circadian control while the CNS sites studied had less than 4% rhythmic genes^7^. Similar low levels of cycling gene expression have been found in the striatum (5%) ^8^ and forebrain cortex (6%)^9^. Ultradian gene expression was first comprehensively described in the mouse liver, revealing a small subset of genes with periods of either 8- or 12-hours^10^. More recent analysis found that 6% of transcripts in the human prefrontal cortex have a 12-hour period, which can be altered in neurological disease^11^. Previous work has observed infradian rhythms among genes, often exemplified by variations in menstrual and seasonal cycles^6^. However, there are few studies of gene rhythms for periods between 24- and 48-hours in the CNS or other organs/tissues. Beyond needing to elucidate the dynamics of cycling gene tissue specificity and variable cycling periods in the naïve CNS, post-transcriptional regulation of these genes adds yet another complex variable to consider in CNS gene expression.

Circadian proteomics studies identify more circadian rhythmicity than what is observed at the transcriptome level in the brain and liver^9, 12^, suggesting that post-transcriptional/translational regulators generate rhythmicity; miRNAs are key post-transcriptional regulators that may contribute to this phenomenon. These short RNAs typically bind to the 3’ untranslated region of mRNAs, leading to decreased mRNA stability and protein production. Through this function, miRNAs play important roles in neuroplasticity and neurodegenerative diseases^13^. Moreover, these RNAs have cycling rhythms of expression across the mouse transcriptome^14, 15^. Potentially due to their own cyclic nature, miRNAs are suggested to modify the rhythmic characteristics of mRNAs^14, 16^. Furthermore, miRNAs act in a highly tissue-specific manner, which may explain the aforementioned tissue-specificity of circadian gene expression^17^.

We, therefore, hypothesized that CNS rhythmicity would occur differentially across regions and cell types and that cycling genes would be co-expressed. Our work identifies site-specific cycling genes in the CNS and works towards uncovering their impact in the naïve state. Potential interactions between cycling mRNAs and miRNAs, whose post-transcriptional regulation of rhythmic genes may be tissue-specific, are also revealed. Finally, we describe clusters of cycling genes and their approximate cellular composition. We validate our findings using cell-specific protein expression. Our results build a foundation upon which potential molecular/cellular circadian gene networks and interactions can be identified across the CNS, which we now provide as a searchable database.

## Results

### Distinct biological rhythms of mRNAs and miRNAs across CNS sites

We began by identifying cyclic mRNAs and miRNAs from tissues collected every 3 hours over 36-hours in the naïve mouse cerebral cortex, hypothalamus, and striatum, with the liver used as a positive control (**Supplementary Fig. 1; Methods**). Bioinformatic guidelines^18^ to detect circadian/rhythmic gene expression were employed for this analysis using the R packages *MetaCycle*^19^, to detect genes with a 24-hour period, and *RAIN*^20^, to detect harmonic (i.e. non-24-hour) rhythms.

Surprisingly, the cortex and not the hypothalamus had the highest number of cycling mRNAs among CNS tissues (**Fig. 1a**; **Supplementary Table 1**). As expected^7^, the liver had the most cycling mRNAs overall, although the cortex had more when considering non-circadian mRNAs. To confirm our results, we compared cycling genes detected by *MetaCycle* and *RAIN*; all genes identified using *MetaCycle* were also found using *RAIN* (**Fig. 1b**). Moreover, the periods for these overlapping genes were largely estimated to be 24±3-hours by *RAIN*, showing concordance between these algorithms (**Fig. 1c**). There was a 1.8 to >5-fold increase in the number of cycling genes identified among CNS sites (with only a ∼20% change in the liver) when using *RAIN*, suggesting the increased presence of harmonic genes in the CNS. Using *MetaCycle*, only 8 cycling miRNAs were identified in at least one site (miR-5099, miR-124a-2, miR-342, miR-150, miR-29b-1, miR16-1, miR-6539, and miR-223), consistent with previous studies showing few cycling miRNAs^7, 14, 21, 22^. As with our mRNA analysis, there were more cycling miRNAs identified using *RAIN*. Hence, the proportions of genes changing across CNS sites align with previous studies^7–9, 19, 23^. We now identify new cycling genes in the CNS due to our use of a broader range of cycling periods and improved data resolution.

**Fig. 1:**
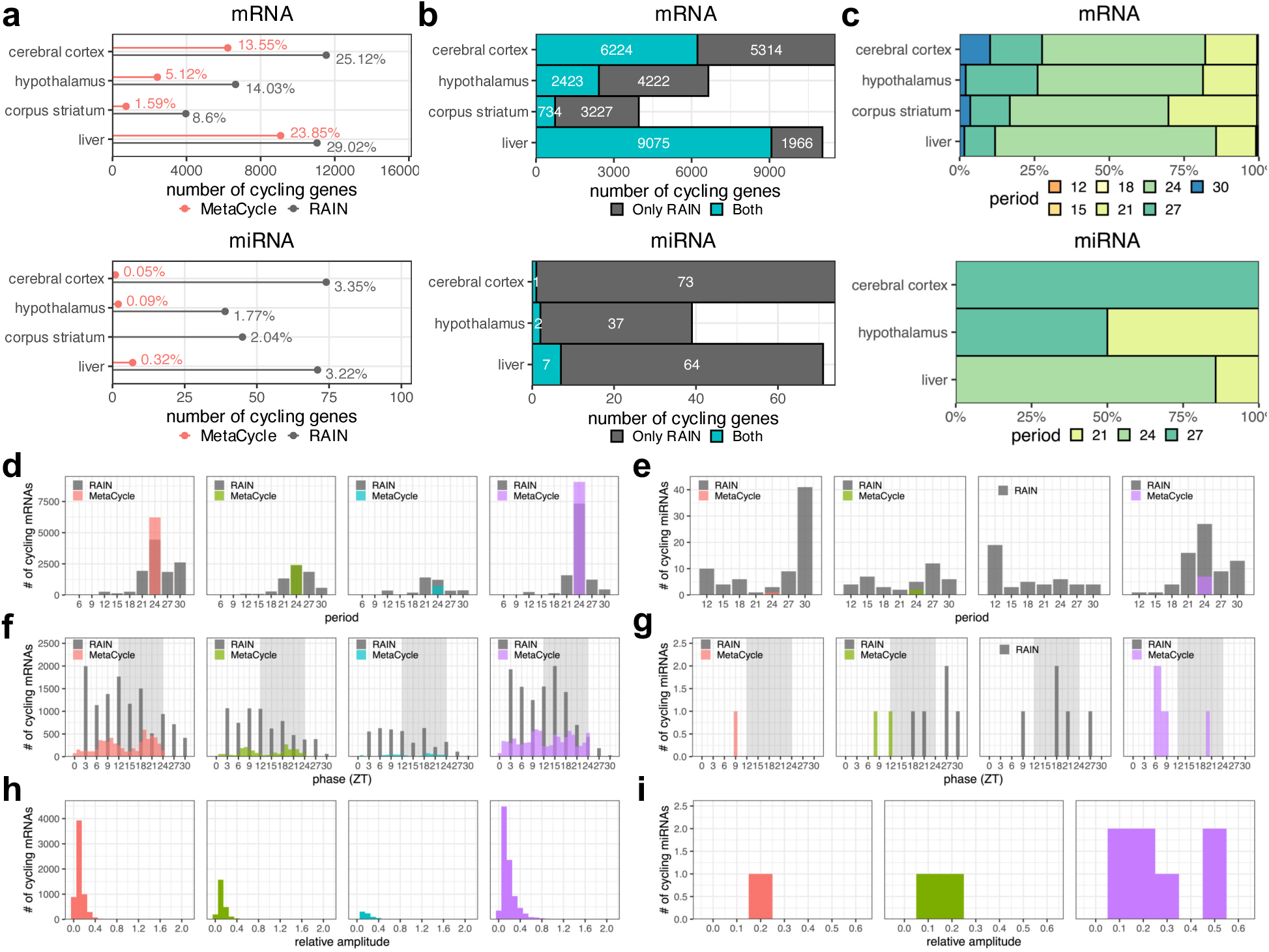
Variability in the number, period, phase, and relative amplitude of cycling mRNAs and miRNAs across CNS tissues. **a**, The number (x-axis; *P*BH<0.05) and percentage (listed in plots) of cycling genes in each CNS tissue, with results using both *MetaCycle* (red) and *RAIN (*grey). **b,** Overlap of the numbers of genes detected by only *RAIN* (grey) or both *MetaCycle* and *RAIN* (turquoise). **c,** The distribution of period lengths (hours) predicted by *RAIN* amongst cycling genes also detected by *MetaCycle*. **d-i,** Cycling parameters estimated in the cerebral cortex (red), hypothalamus (green), corpus striatum (turquoise), and liver (purple) using *MetaCycle* and *RAIN* (grey). **d-e,** Distribution of period lengths between 6 and 30-hours. **f-g,** Distribution of phases between Zeitgeber Time 3 and 30, with entrained darkness highlighted with a grey background **h-i,** The distribution of relative amplitudes from 0.015 to 2.03.

As aforementioned, most cycling genes appear circadian, though ultradian and infradian periods are also present in the CNS. As shown by others^10, 19^, we found the period of liver mRNAs clustered around 12-, 24-, and >24-hours (**Fig. 1d**; **Extended Data Figs. 1-3**). While these patterns also appear to a lesser extent in the CNS, most genes exhibited a 24±3-hour period (**Fig. 1d,e**). Across all tissues, mRNA and miRNA phases peaked during the light/dark transition (**Fig.1f,g**). Relative amplitudes had a wide range across tissues, with the liver having the greatest range whereas the CNS was more restricted (**Fig.1h,i**). We focused on the expression of key clock genes (**Extended Data Fig. 4**), and confirmed our expectation that these genes would have a 24-hour period. Among core clock genes, the transcription factor *Bhlhe41* was only rhythmic across the CNS*, Adrb1* was only cycling in the cortex, and *Timeless* was only rhythmic in the liver.

### CNS specificity of cycling genes

We then assessed the site-specificity of detected cycling genes by focusing on the more comprehensive results obtained using *RAIN*. Relatively few cycling mRNAs (n=1332) and miRNAs (n=1, miR-5099) were detected across all CNS sites and the liver (**Fig. 2a,b**), with the liver containing the most unique cycling genes and the striatum the fewest. When excluding the liver from this analysis and focusing only on cycling genes shared across CNS sites, a greater number of cycling mRNAs (n=2113) were identified, with again only miR-5099 showing rhythmicity among miRNAs. Moreover, many of these shared genes exhibited different cycling parameters (i.e., period) depending on the site assessed (**Supplementary Table 2**). Period differences were identified in 7,391 mRNAs and ranged between 3- and 21-hours across sites (**Fig. 2c,d**). We identified 7,235 mRNAs with phase differences between any two CNS sites, of which 3,955 had an absolute difference ≥6-hours, including 1,131 in antiphase (12±3-h; **Fig. 2e,f**). *MetaCycle* was used to measure relative amplitudes, a feature not available in *RAIN*, and identified 495 shared CNS genes with relative amplitudes up to 0.71 (**Fig. 2g**).

**Fig. 2:**
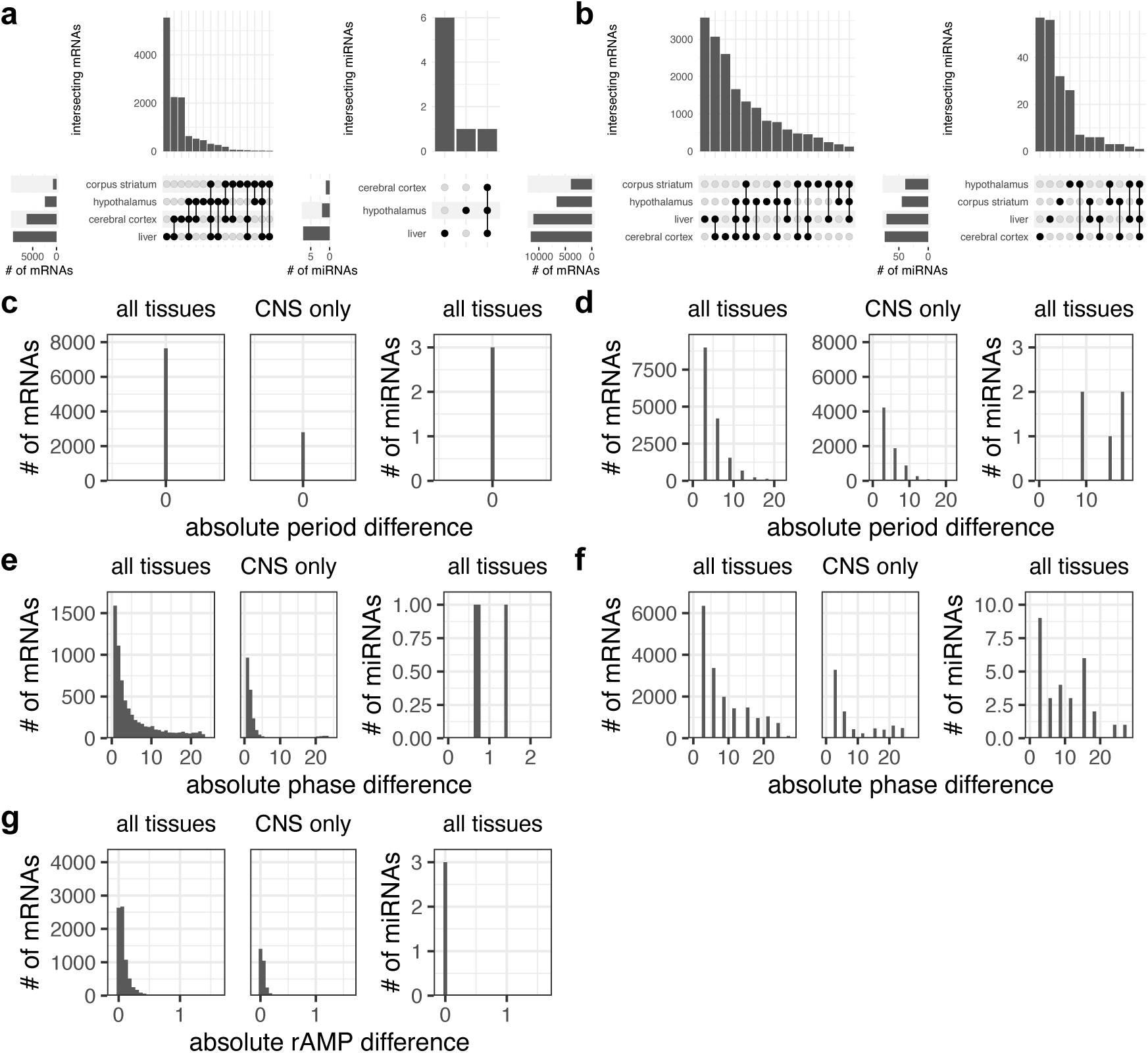
Comparison of cycling genes across tissues. **a,b**, Intersections of cycling genes **a,** Comparison of cycling genes from *MetaCycle* across tissues. mRNAs are in the left panel, and miRNAs are in the right. **b,** Comparison of cycling genes from *RAIN* across tissues. mRNAs are in the left panel, and miRNAs are in the right. **c-g,** Distributions of cycling parameter differences between tissues. **c,e,g,** Cycling parameters estimated by *MetaCycle*. **d,f,** Cycling parameters estimated by *RAIN*.

### Cycling genes are linked to human chronotype genetics

Given that the cycling mRNAs identified by *MetaCycle* are estimated to have a 24-hour period, and thus may be under circadian control, we assessed their overlap with 366 genes previously linked to human chronotype^24^. Of these, cycling genes were identified in the cortex (n=107_, hypothalamus (n=39), striatum (n=10_, and liver (n=128_. Only *Rbm6*, *Fbxl4*, *Rfx4*, *Ahsa2*, and *Hexim1* were shared across all CNS sites (**Supplementary Table 3**). Transcription corepressor binding, which includes genes that repress gene expression by binding to transcription factors, and circadian rhythm/entrainment were key pathways identified among chronotype genes; no pathways were shared across all CNS sites (**Fig. 3**; **Supplementary Table 4**). These genes may be critical to chronotype predisposition, which is known to vary across mouse strains^25^.

**Fig. 3:**
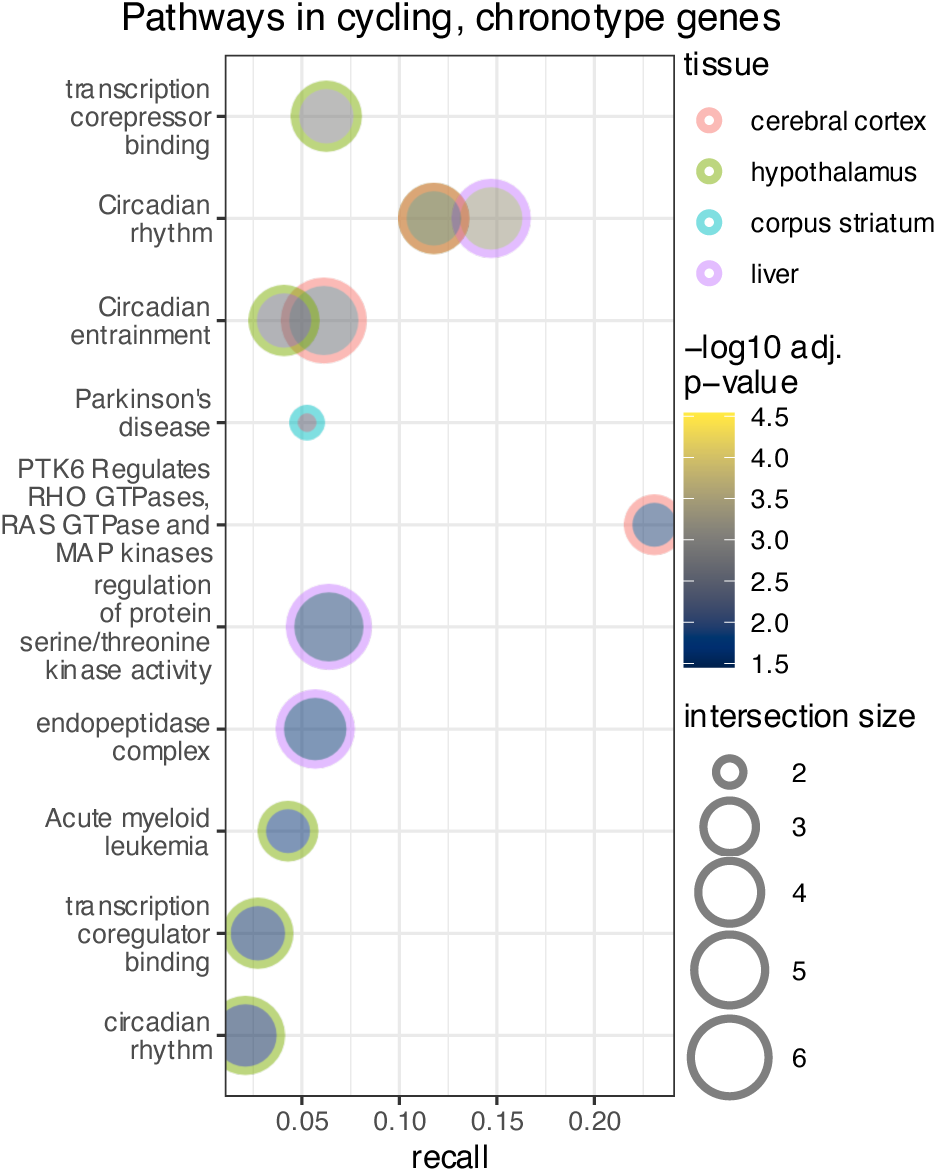
DNA variants linked to chronotype are enriched in circadian and transcription regulation pathways. The top 10 pathways enriched in genes that are both cycling and linked to DNA variants associated with chronotype (*P*_g:SCS_<0.05; 10≤term size≤500). Intersection size represents the number of genes in a pathway that are cycling, and recall represents the proportion of genes represented in each term/pathway.

### Shared and unique pathways across the CNS

Analysis of cycling mRNAs from *RAIN* found protein folding, cell migration, and RNA modification as shared pathways across all tissues (**Fig. 4a**; **Supplementary Table 5**); those unique to the CNS included axonogenesis, learning or memory, mRNA stabilization, and macrophage differentiation (**Fig. 4b-d**; **Supplementary Table 6**), while ribosomal and metabolic pathways were unique to the liver (**Fig. 4e**; **Supplementary Table 7**). Further, we identified pathways that are uniquely enriched in the 12±3, 24±3, and 28-30-hour period categories. Pathways involving autophagy, protein folding, and cellular stress were unique to the mRNAs with a 24±3 period. (**Extended Data Fig. 5a,c,f,i; Supplementary Table 8**). Pathways unique to the 12±3 mRNAs align with existing literature, showing that 12-hour cycling genes in the liver are enriched in the endoplasmic reticulum and Golgi processes (**Extended Data Fig. 5b,g,j**)^26^. The top pathways unique to the 28 to 30 category in the cortex were notably involved in synaptic activity, and those in the liver had a strong innate immunity signature (**Extended Data Fig. 5c,e,h,k**).

**Fig. 4:**
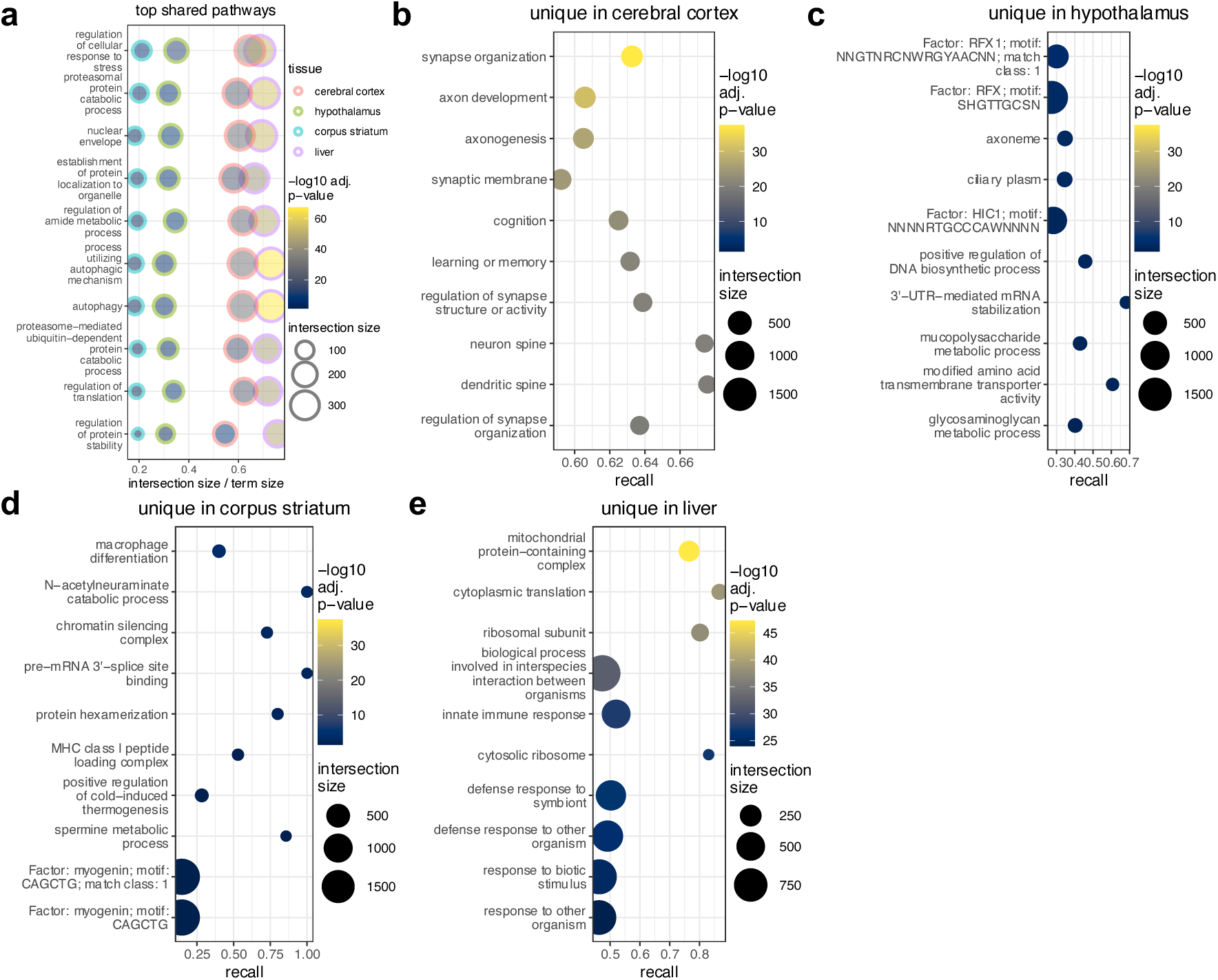
Comparison of pathways enriched in cycling genes across tissues. **a**, Top pathways that are enriched in all 4 tissues (*P*_g:SCS_<0.05; 10≤term size ≤500). Intersection size is the number of genes that are both cycling and in a pathway. Recall is intersection size divided by the size of a term/pathway. **b-e,** Top pathways uniquely enriched in their respective tissues (*P*_g:SCS_<0.05; 10≤term size ≤500).

### Identification of mRNA-miRNA pairs and pathways across the CNS

Associations between cycling mRNA-miRNA pairs in the CNS were tested and yielded 5,618 pairs in the cortex, 2,533 in the hypothalamus, and 42,659 in the liver. Putative direct control of mRNA expression by miRNAs was identified using a negative correlation and delay ≤0 in ∼50% of genes in the CNS and ∼25% in the liver **(Supplementary Tables 9**), suggesting that miRNAs may play a more important role in CNS rhythmicity than in the liver. Of the direct interaction pairs, only a small subset has been previously predicted or experimentally validated (**Fig 5**; **Supplementary Table 10,11**), highlighting the need for further study of miRNA regulation of circadian gene expression. Previously predicted but unvalidated direct mRNA-miRNA cycling pairs may present a starting point for assessing post-transcriptional regulation of CNS rhythmicity.

**Fig. 5:**
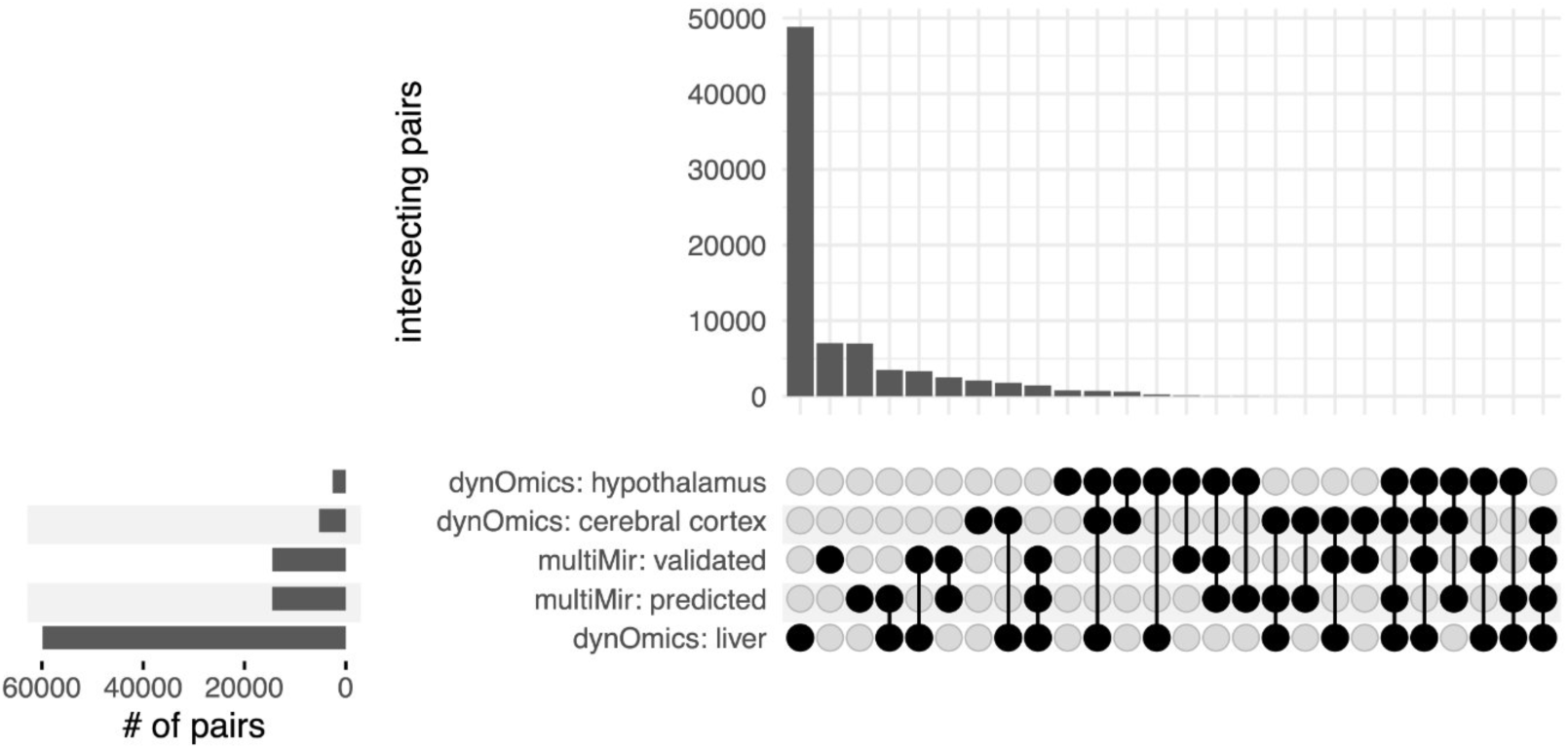
mRNA-miRNA association analysis. Intersections between mRNA-miRNA pairs associated (P_BH_<0.05) across tissues and previously reported pairs from *multiMir*.

Since genes work in concert to affect complex biological functions, we used network analysis to identify co-expression modules of mRNAs and miRNAs separately (**Extended Data Fig. 6,7; Methods**). Cycling mRNAs were enriched in 12 of 26 modules in the cortex, 7 of 15 in the hypothalamus, 9 of 13 in the striatum, and 7 of 20 in the liver (**Fig. 6a,b**; **Supplementary Table 12**); the majority of hub genes in these modules were rhythmic. *RAIN* cycling miRNAs were in all modules except one striatal module; *MetaCycle* miRNAs were in 1 of 4 modules in the cortex, 2 of 3 in the hypothalamus, and 3 of 3 in the liver (**Fig. 6c,d**). By visualizing the composition of cycling genes and their period categories within modules, we found that most modules contain genes with a period 24±3-hours (**Extended Data Fig. 8,9**). However, there are some exceptions; for example, the cortical *yellow4* module has very few non-cycling mRNAs, of which most have a period of 27-hours or greater. Finally, analysis of cycling module eigengenes showed that 33 of 126 mRNA-miRNA module pairs were associated (P_BH_<0.05; **Extended Data Fig. 10**). Altogether, our study identifies modules of coordinated rhythmic gene expression in the CNS.

**Fig. 6:**
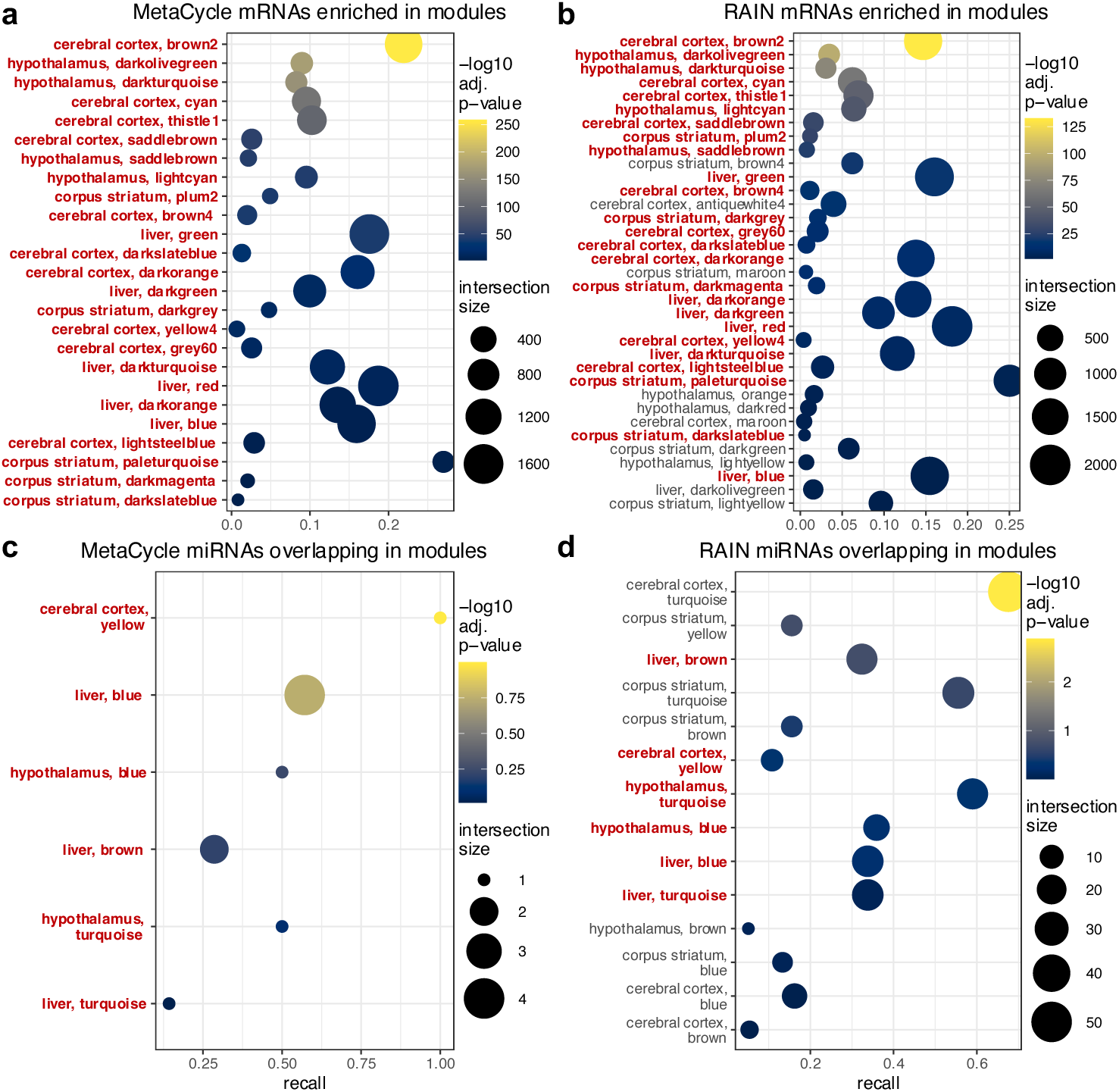
Network analysis and enrichment of cycling genes. **a-d**, Overlap between gene modules and cycling genes. The sizes of datapoints represent the number of genes that are both cycling and in a module, called intersection size. Recall is the intersection size divided by the total number of cycling genes considered. Datapoints are coloured by the -log10(*P*_PH_) of cycling gene enrichment in a module. **a,b,** mRNA modules enriched with cycling genes (*P*_BH_<0.05). **c,d,** miRNA modules that contain at least one cycling gene. **a,c,** Analysis of *MetaCycle* cycling genes in modules. **b,d,** Analysis of *RAIN* cycling genes in modules. Modules that occur in both *MetaCycle* and *RAIN* results are emphasized by bold red text.

### Chronobiological rhythms contribute to neuroimmune interactions in the healthy state

The CNS is a complex organ including neuronal, glial, and supporting cells. Thus, we examined the cellular composition of cycling mRNA modules that are typically thought to represent distinct cell types, although recent evidence shows heterogeneous modules in the striatum^23^. CNS cell marker genes were enriched in 8 cycling modules in the cortex, 1 in the hypothalamus, and 3 in the striatum; as expected, none were enriched in the liver (**Fig. 7a**; **Supplementary Table 13**). Most modules appear heterogeneous and contain gene markers for multiple glial/immune cells, such as oligodendrocytes, microglia/perivascular macrophages, and astrocytes. For example, the cortical *grey60* module (enriched in oligodendrocyte, astrocyte, and endothelial cell markers) is enriched in pathways including myelination, glial cell differentiation, and oligodendrocyte differentiation (**Fig. 7b,c**; **Supplementary Table 14**). These results suggest that rhythmic cell-cell communication may play an important role in driving CNS biology.

**Fig. 7:**
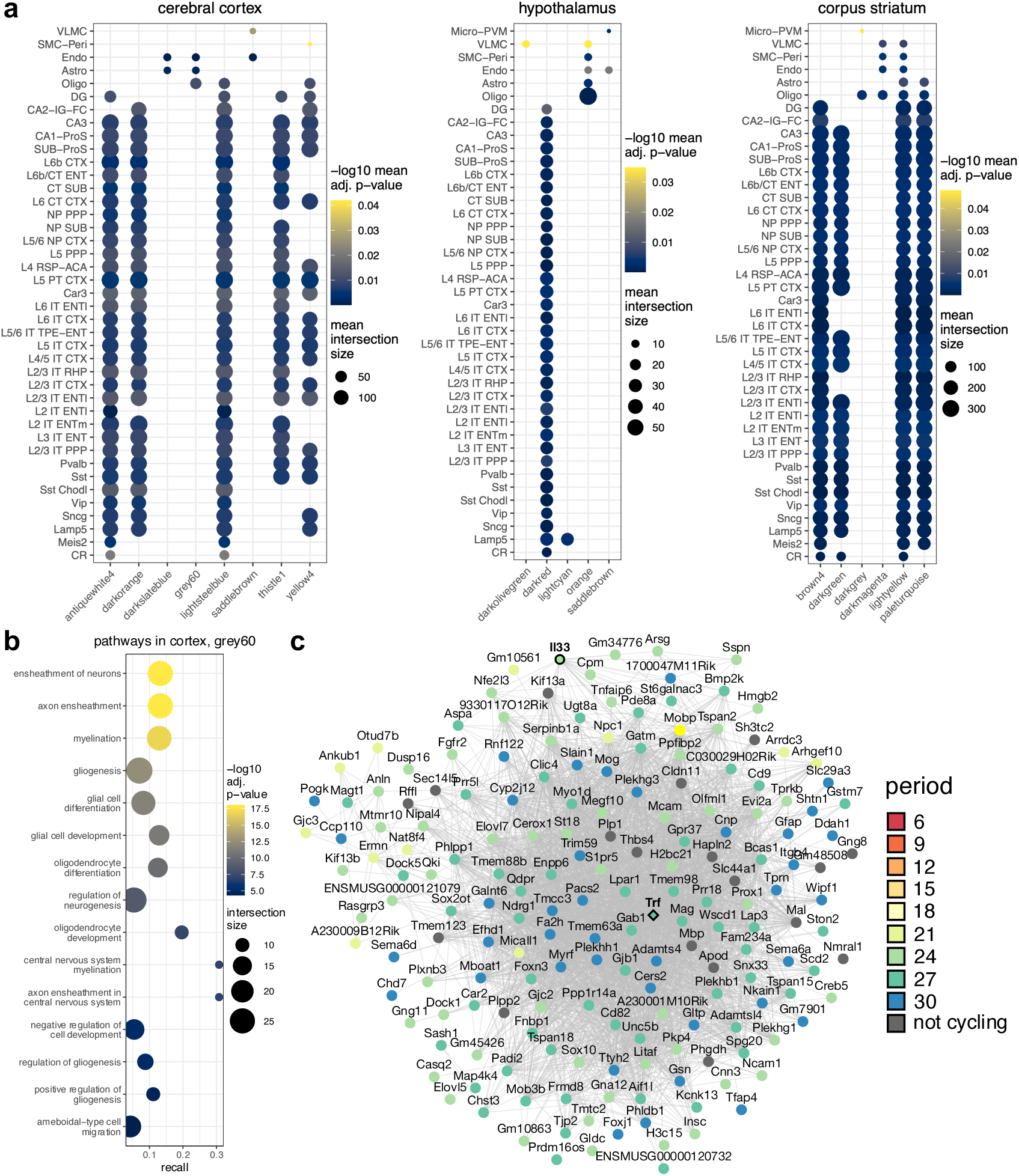
Enrichment of cell type marker genes in cycling mRNA modules. **a**, Cell type clusters (n=387) are collapsed into 42 previously defined subclasses for visualization. *RAIN* cycling mRNA modules enriched in cell type marker genes and their respective cell type subclasses are shown (*P*_BH_<0.05). The size of datapoints represents the mean number of genes that are both a cell subclass marker gene and in a module, i.e., mean intersection size. Datapoints are coloured by the mean -log10(*P*_PH_) of a cell type subclass’s marker genes enrichment in a cycling module. **b,** Pathways enriched in the cortical *grey60* module (*P*_g:SCS_<0.05; 10≤term size≤500). **c,** mRNAs with an adjacency≥0.03 are visualized (n=180 of 290). mRNAs (circle nodes) are arranged based on co-expression adjacency (grey edges). Nodes of cycling genes are coloured by period category. If a gene is not cycling, its node is coloured grey. The hub gene has a diamond node. Genes that underwent immunohistochemistry analysis, *Trf* and *Il33*, have their nodes outlined in black and labels bolded.

Finally, we sought to detail the cell- and site-specific nature of cycling genes at the protein level and focused on the *grey60* module in the cortex, given its specificity for glial cells. The hub gene in this module, transferrin, has previously been shown to be expressed by oligodendrocytes^27^ and showed rhythmic expression with a trough at ZT2 and a peak at ZT14 (**Fig. 8a,b**). IL-33, also in this module and known to play a role in myelination^28, 29^, displayed circadian changes in immunofluorescent intensity in cortical Olig2+ cells that peaked at ZT14 (**Fig. 8c,d**; **Extended Data Fig. 11a,b, 12a**). These results align with recent data showing that oligodendrocyte precursor cell (OPC) dynamics are subject to time-of-day differences in the naïve cortex via *Bmal1*, regulating sleep architecture and OPC complexity^30^. Candidate genes *Fzd4*, *Eif1b*, *Lrrk2*, *Sphk2*, and *Adipor2* were also selected for protein analysis given their strong rhythmicity and differing phases in the cortex and hypothalamus. We found these targets to be expressed rhythmically at the protein level in neurons across the CNS. AdipoR2 showed rhythmic expression in hypothalamic neurons (peaking at ZT14), while Fz-4 expression peaked at ZT8 in cortical neurons (**Extended Data Fig. 11a,c,d, 12b,c**). Eif1b, on the other hand, exhibited rhythmic immunofluorescence in cortical neurons with increased expression at ZT20/2 and reduced expression at ZT8/14 (**Extended Data Fig. 11a,e, 12d**). Sphk2 and Lrrk2 co-localized with neurons in the cortex but had no significant changes in fluorescence intensity across the timepoints examined (**Extended Data Fig. 11a,f,g, 12e,f**). These findings highlight the need for cataloguing changes in rhythmic expression at both the gene- and protein-level, in a cell- and site-specific manner.

**Fig. 8:**
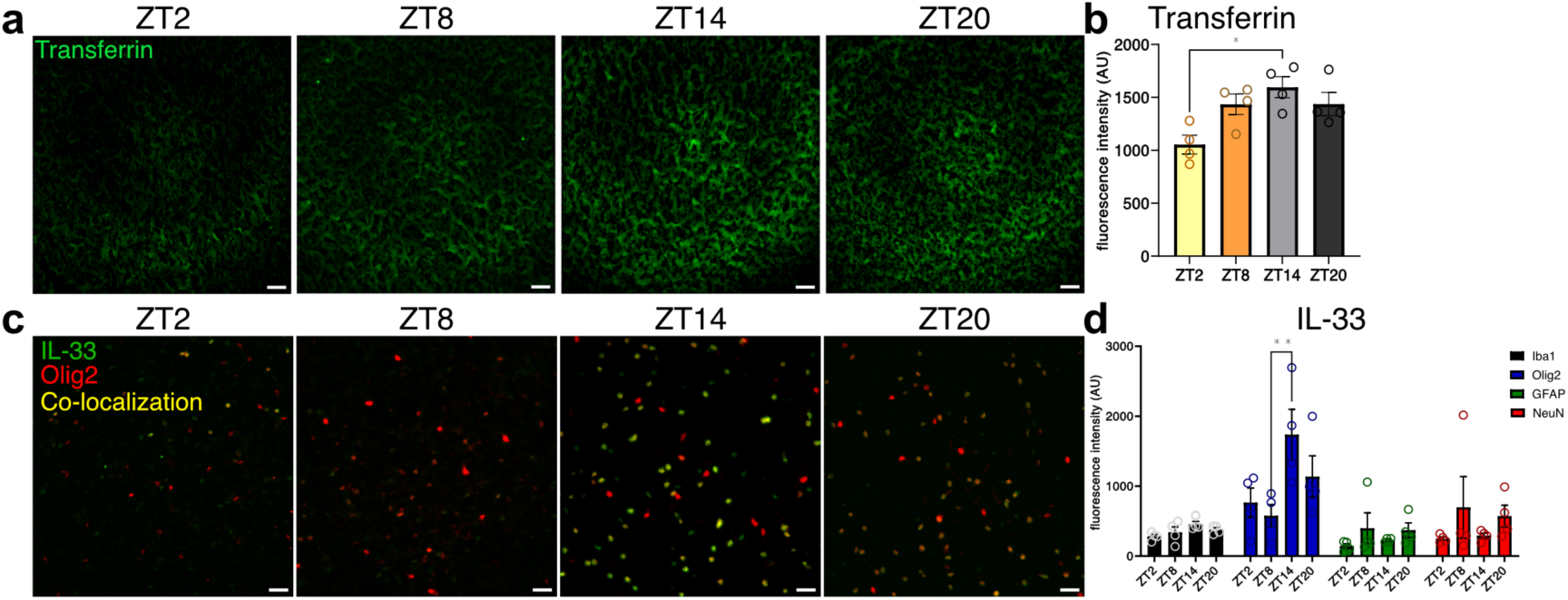
Rhythmic expression of extracellular transferrin and oligodendrocyte-specific IL-33 in the naïve CNS. **a**, Representative images of transferrin staining in the cortex of brains collected at ZT2, ZT8, ZT14, and ZT20. Scale bar, 25µm. **b,** Quantification of fluorescence intensity of transferrin signal in the cortex. One-way ANOVA with Tukey’s post hoc test, *n*=4 animals for each timepoint. Transferrin ZT2 versus ZT14, \**P*=0.0106, *q*=5.457, d.f.=12. **c,** Representative images of cortical oligodendrocytes (labelled with Olig2), IL-33, and their co-localization at ZT2, ZT8, ZT14, and ZT20. Scale bar, 25µm. **d,** Quantification of fluorescence intensity of IL-33 signal co-localized in the cortex with Iba1 (microglia), Olig2 (oligodendrocyte marker), GFAP (astrocytes), and NeuN (neurons). Two-way ANOVA, with Tukey’s post hoc test, n=4 animals for each timepoint. IL-33 with Olig2, ZT8 versus ZT14, \*\**P*=0.0070, *q*=6.093, d.f.=48. All data presented as mean ± s.e.m.

## Discussion

We identify cycling mRNAs, miRNAs, gene networks, and novel mRNA-miRNA co-expression pairs in the naïve CNS, which were validated at the protein-level with cellular resolution. Importantly, these rhythmic changes were both tissue- and cell-specific, highlighting the different expression characteristics of cycling genes and their variability across CNS sites. Thus, circadian rhythmicity must be considered when planning experiments assessing gene function/activity and behavioural outcomes, for which our curated catalogue can be a starting point (https://www.ghasemloulab.ca/chronoCNS/).

Our study confirms previous findings that circadian gene expression is highly tissue-specific, especially since cycling genes shared across tissues may have different parameters (e.g., phase)^7^. Further, the known function of these genes is also tissue-specific. For example, cycling genes in the cortex are uniquely involved in synapse organization and axonogenesis. This result is striking since that the study that found that 6% of the forebrain cortex’s transcriptome is cycling also found that synaptic transcripts are under strong circadian control^9^. The same group also found that there were more cycling synaptic proteins than synaptic transcripts, meaning that rhythmicity was most likely set at the post-transcriptional stage. To our knowledge, this study is the first to identify putative miRNA-mRNA pairs that could explain this discrepancy. These results have important translational potential given that both circadian rhythms and miRNAs have roles in neurodegenerative diseases, in which synaptic plasticity is disrupted, such as Alzheimer’s Disease and Parkinson’s Disease^31, 32^. For instance, several studies now show the utility of circadian miRNAs (or, circaMiRs) as biomarkers^33–35^ in both healthy and disease states.

We identified groups of co-expressed mRNAs and miRNAs using gene network analysis, many of which were over-represented by cycling genes, with some having signatures of neuronal and/or glial cells. For example, the cortical *grey60* module was only enriched in glial cell markers and gliogenesis pathways. In this module, *Il33* exhibited rhythmic protein expression in oligodendrocytes. Meanwhile, the hypothalamus-specific *saddlebrown* cluster was only enriched in neuronal markers, with AdipoR2 being rhythmic particularly in neurons. These results confirm evidence from others that cycling gene expression is cell-specific^36, 37^, and further support evidence that both glia and neurons are under circadian control^5, 37^. Disruption of these rhythms likely contributes to clinical pathologies including chronic pain, neurodegenerative and demyelinating diseases^38, 39^.

The transcriptomic rhythmicity of these sites has previously been described and, in addition to other tissues, can be explored using various databases, notably CircaDB^40^, CGDB^41^, CircadiOmics^41^, and CircadiOmics^42^. However, these databases either rely on microarrays and not the more robust RNA-sequencing, fail to account for site-specific rhythms, do not capture an adequate number of data points to account for ultra/infradian rhythms, and/or lack data on miRNAs. And yet, it is likely that rhythms are governed at the cellular level. Single-cell circadian datasets of the CNS are only available for the murine suprachiasmatic nucleus and liver, and the much less complex *Drosophila* brain^36, 37, 43^. A recent spatial cell-type atlas of the mouse brain contains samples taken from both the light and dark phase^5^. While this dataset captured diurnal changes of clock genes, non-clock genes were not assessed. As such, more comprehensive databases using single-cell methods are necessary to fully capture the extent of rhythmicity at both site- and cellular-specific levels.

We provide the first examination of mRNA and miRNA rhythmicity, and their interactions, in the mouse cortex, hypothalamus, and corpus striatum. Our work builds a foundation for the study of circadian rhythms in the CNS by better understanding the naïve state, particularly given the contribution of identified cycling genes across neurological diseases including Parkinson’s^44, 45^, stroke^46^, and glioma^47, 48^, among others.

## Methods

### Transcriptomics analysis study design

Our curated workflow for transcriptomics data analysis is summarized in **Supplementary Fig. 1** and detailed in the following sections.

**Supplementary Fig. 1:**
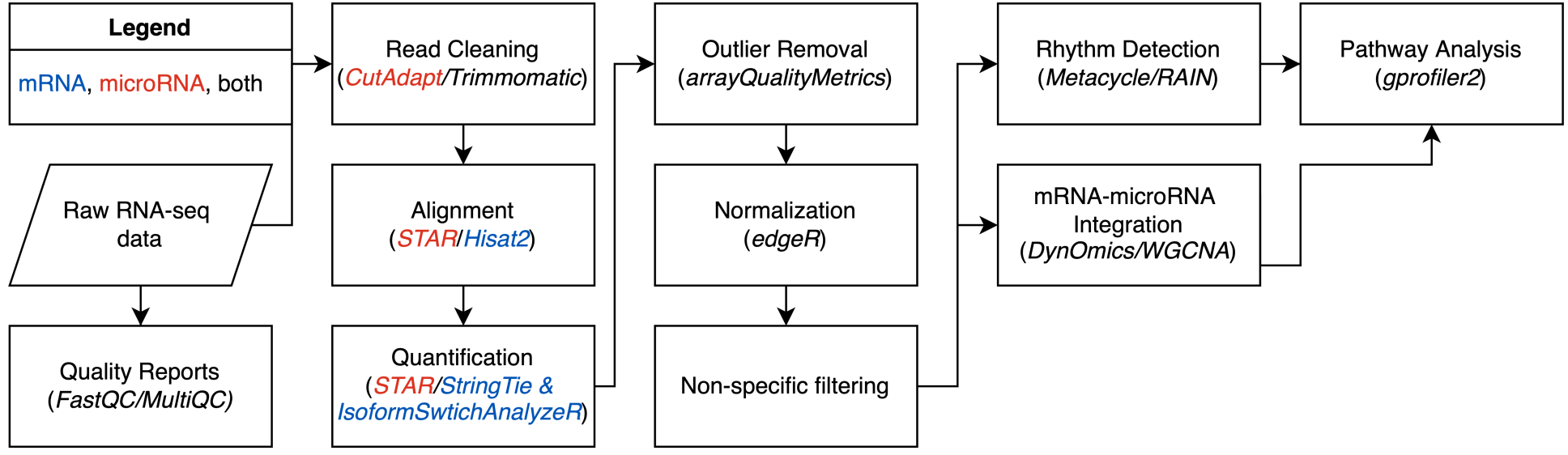
Flow chart illustrating the analysis workflow. Key tools used to perform each step are in italics. First, sequencing quality reports were generated using *FastQC* and *MultiQC*. miRNA reads’ 3’ adapters were removed with *CutAdapt*. Low-quality base pairs were filtered from mRNA and miRNA reads. Then, *STAR* aligned miRNA reads to the mouse reference genome and quantified gene abundance. mRNA reads were aligned to the mouse genome using *Hisat2*, transcript abundance was calculated with *StringTie,* and gene count estimation was performed with *IsoformSwitchAnalyzeR*. On mRNA and miRNA gene counts, we then performed sample outlier removal, normalization, and non-specific filtering. *Metacycle* and *RAIN* were used to detect cycling genes. *DynOmics* detected pairs of mRNA-miRNA cycling pairs. *WGCNA* constructed networks of co-expressed mRNAs and co-expressed miRNAs, whose rhythmicity and association were tested. Finally, pathway analysis of cycling genes, mRNA-miRNA pairs, and co-expressed genes was performed with *gprofiler2*. See Methods for more details.

### Transcriptomics data description

We used *SRA-Toolkit* v2.10.8 to download data from Gene Expression Omnibus (GEO): GSE151567^49^. Morton and colleagues collected samples from the cerebral cortex, corpus striatum, hypothalamus, and liver of 26-week-old wild-type C57BL/6J male mice every 3-hours for 36-hours. Time 0 represents the beginning of daylight, and 12 is the beginning of darkness. After tissues were harvested, they were flash-frozen and transferred to Expression Analysis, Inc. for mRNA and miRNA extraction and sequencing. There were 2-6 biological replicates for each timepoint, with the exception that there were no miRNA-seq samples from Zeitgeber time 15 in the striatum (**Supplementary Fig. 2a**). Data collection is further described by Wang *et al*^23^.

**Supplementary Fig. 2:**
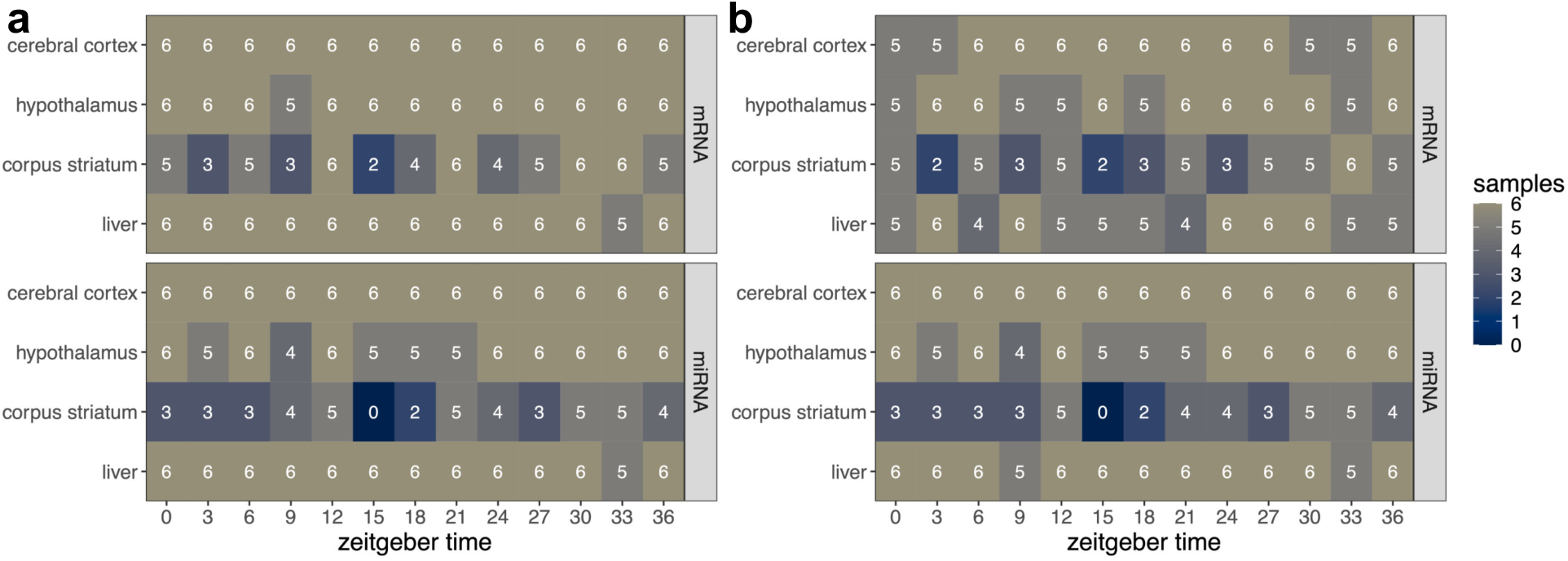
Study design before and after sample exclusion. **a**, Number of samples per timepoint and tissue downloaded from Gene Expression Omnibus (GEO): GSE151567. **b,** Number of samples per timepoint and tissue after quality control of samples.

### Transcriptomics data preparation

We evaluated the quality of sequencing reads with *FastQC* v0.11.9 and *MultiQC* v1.12^50, 51^. For mRNA-seq reads, we filtered for base quality with *Trimmomatic* v0.36 and aligned reads to the mouse genome (GENCODE release M32) with *Hisat2* v2.2.1^52–54^.For mRNA-seq reads, we filtered for base quality with *Trimmomatic* v0.36 and aligned reads to the mouse genome (GENCODE release M32) with *Hisat2* v2.2. Gene expression was quantified with *StringTie* v2.1.5, and R package *IsoformSwitchAnalyzeR* v1.18.0^55, 56^. Adapted from Rahmanian *et al*.’s miRNA-seq processing pipeline, we removed 3’ adapters from miRNA reads, filtered for base quality, aligned, and quantified gene expression using *CutAdapt* v4.2, *Trimmomatic* v0.36, and *STAR* v2.7.9a^52, 57–59^. Overall and unique alignment rates were calculated to evaluate performance. One mRNA and one miRNA sample from the striatum were removed before quantification due to low overall alignment rates (35% and 34.66%, respectively; **Supplementary Fig. 3**).

**Supplementary Fig 3.**
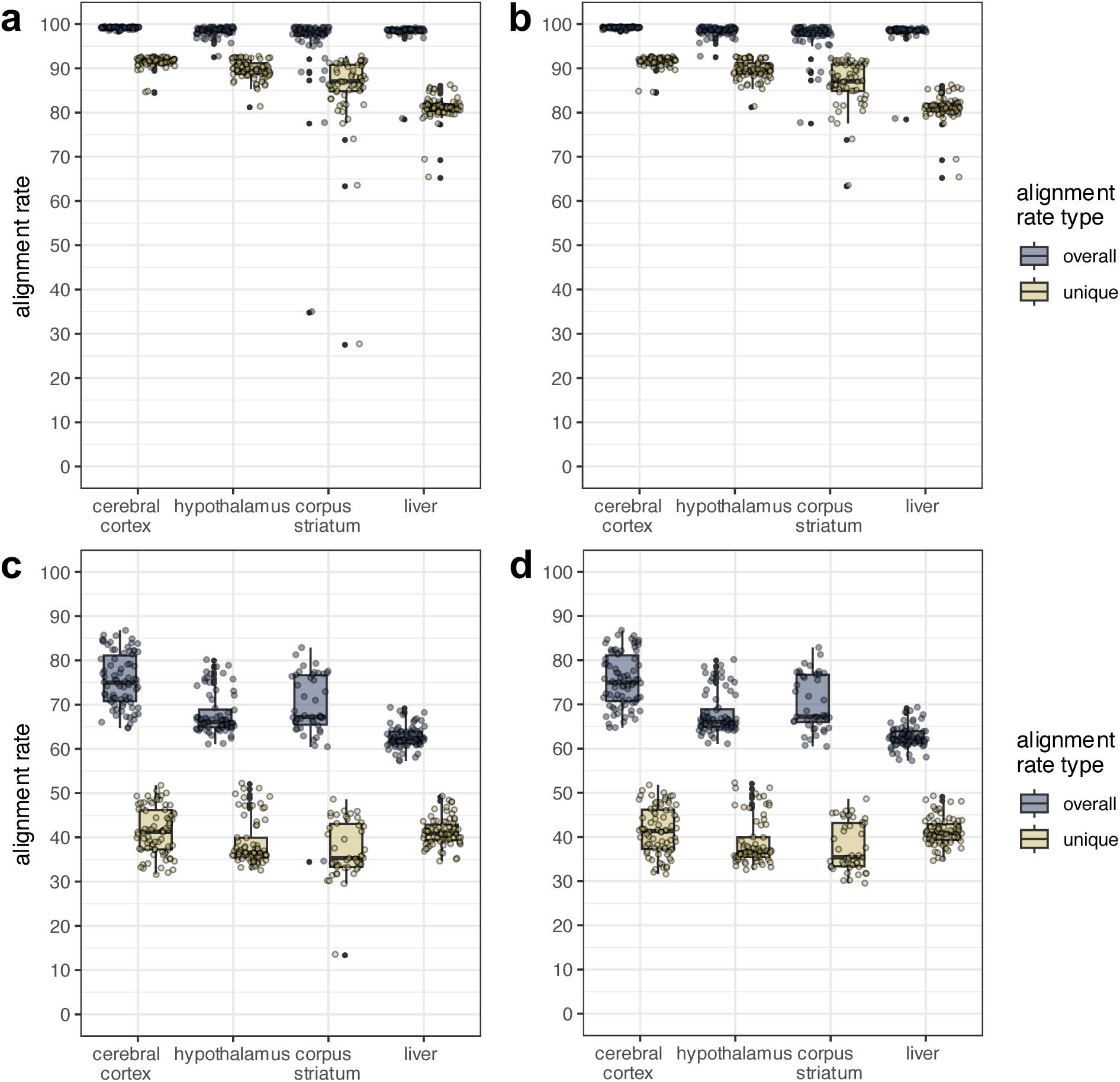
Alignment rates for mRNA and miRNA datasets. The overall alignment rate is the number of RNA reads that are mapped at least once to the reference genome divided by the number of reads in the dataset. The unique alignment rate is the number of reads that are aligned to exactly one location on the reference genome. Boxplots show median and interquartile range (IQR; box size). The maximum and minimum whiskers represent Q3 + 1.5 × IQR and Q1 − 1.5 × IQR. IQR, interquartile range. a,b, Alignment rates for the mRNA dataset. a, Rates before sample SRR11902313 from the striatum was removed due to a low overall alignment rate (35%). b, Rates after the outlier sample was removed. c,d, Alignment rates for the miRNA dataset. c, Rates before SRR11902764 from the striatum was removed due to a low overall alignment rate (34.66%). d, Rates after the outlier sample was removed.

With the generated gene count matrixes, we evaluated this experiment’s technical variability by finding the Spearman correlation between samples from the same tissue and timepoint (**Supplementary Fig. 4**). Then, we performed outlier detection and TMM normalization using the R packages *arrayQualityMetrics* v3.60.0 and *edgeR* v4.2.0, respectively^60, 61^. *arrayQualityMetrics* uses three metrics to consider a sample an outlier: 1) its sum of distances to other samples, 2) the Kolmogorov-Smirnov statistic, and 3) the Hoeffding’s *D*-statistic. Samples were removed if marked an outlier before and after normalization, or if multiple metrics marked them an outlier after normalization. After removing outliers, we re-normalized gene counts, and outlier detection was repeated. Based on these metrics, we removed four (cortex), two (hypothalamus), five (striatum), and nine (liver) samples from the mRNA dataset. One striatum and one liver sample were removed from the miRNA dataset. We used principal component analysis on variance stabilizing transformed counts to visualize the clustering of samples with *DESeq2* v1.44.0 (**Supplementary Fig. 5**)^62^. Two mRNA hypothalamus samples appeared to be outliers and were removed. The final study design is displayed in **Supplementary Fig. 2b**. Finally, we considered genes to be expressed for each tissue if their mean normalized counts per million were >= one. There were 18,444 (cortex), 18,712 (hypothalamus), 18,248 (striatum), and 13,514 (liver) mRNA genes remaining. For the miRNA dataset, there were 459 (cortex), 462 (hypothalamus), 458 (striatum), and 284 (liver) remaining miRNAs.

**Supplementary Fig 4.**
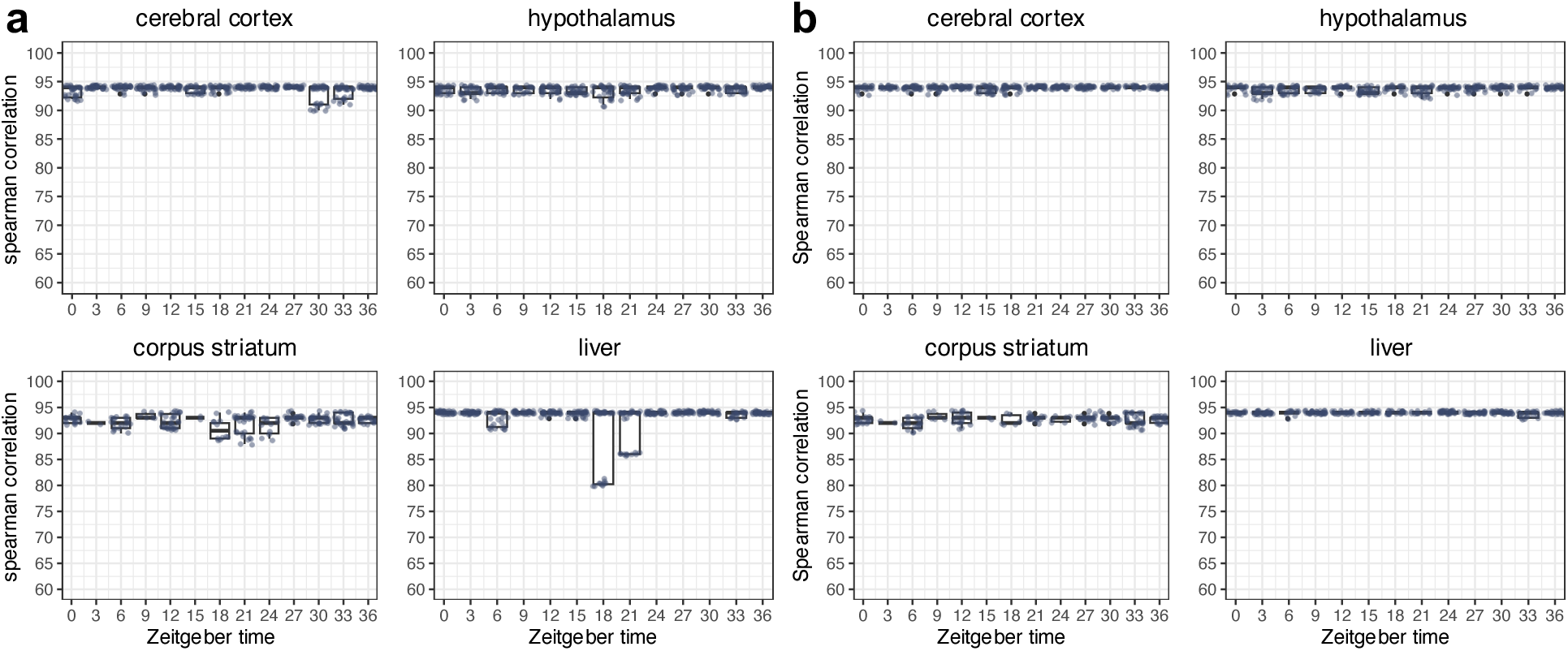
Spearman correlation between samples from the same tissue and sampling timepoint. Sampling timepoints represented in Zeitgeber time are on the x-axis. The Spearman correlation coefficients between replicates are on the y-axis. **a,** Correlation coefficients before outlier removal. **b,** Correlation coefficients after outlier removal. Boxplots show median and interquartile range (IQR; box size). The maximum and minimum whiskers represent Q3 + 1.5 × IQR and Q1 − 1.5 × IQR. IQR, interquartile range.

**Supplementary Fig. 5.**
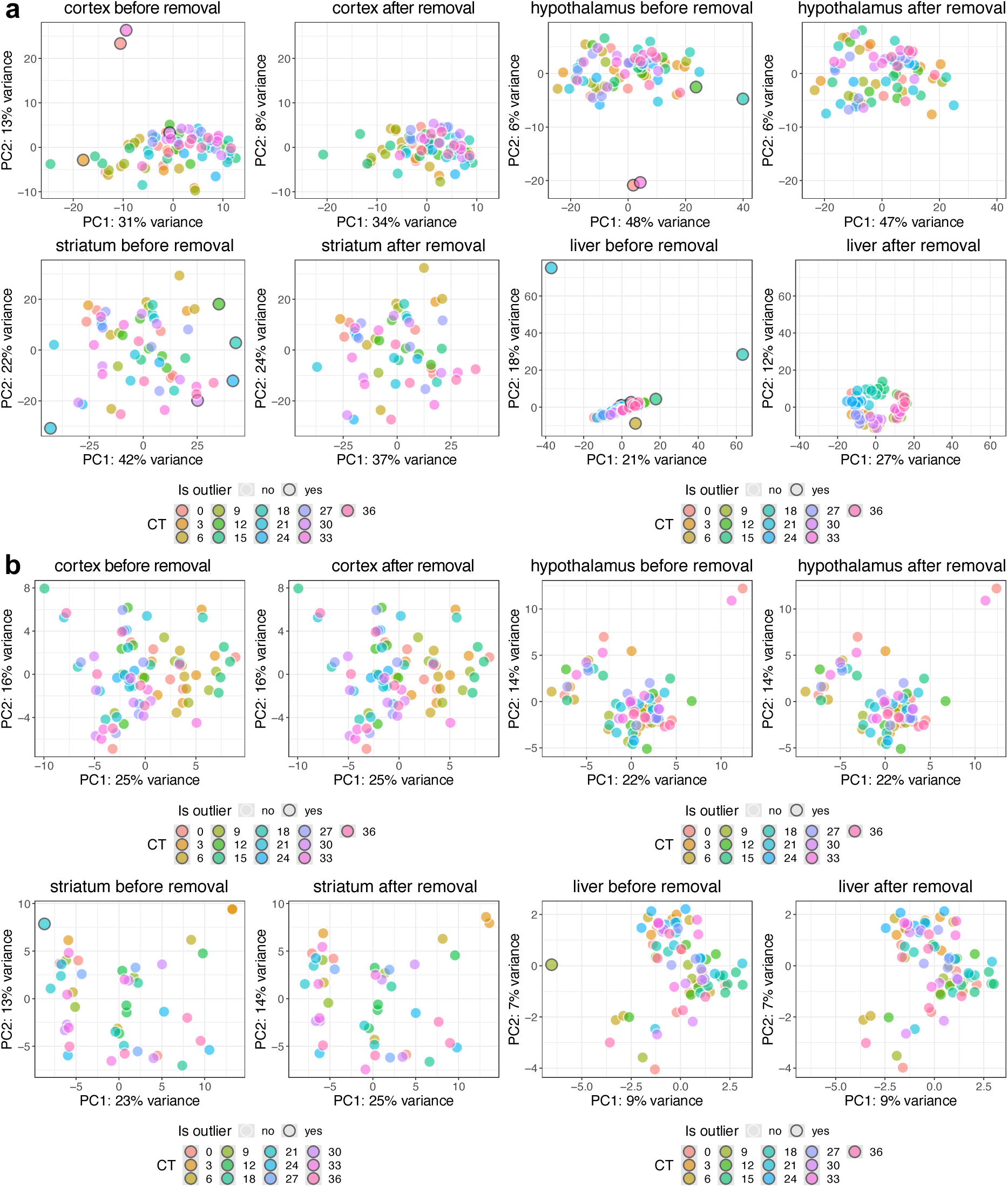
Principal component analysis of samples before and after outlier removal. The x-axes represent the first principal component to explain the variance between samples. The y-axes represent the second principal component. Each data point is a sample. Samples are coloured by timepoint, represented in Zeitgeber time (ZT); samples are outlined in black if they were marked as an outlier and removed. As indicated by titles, panels are grouped by tissue. Within each panel group, the left panel indicates sample clustering before outlier removal, and the right is after outlier removal. **a,** mRNA datasets. **b,** miRNA datasets.

### Identifying rhythmic features

Cycling mRNA and miRNAs were detected using R packages *MetaCycle* v1.2.0 and *RAIN* v1.38.0^19, 20^. For *MetaCycle*, we used the meta2d function with a minimum and maximum period of 24. *P*-values from the Lomb-Scargle (LS) and Jonchkheere-Terpstra-Kendall (JTK) cycle algorithms were combined with Fisher’s method and corrected for multiple testing with the Benjamini-Hochberg method. For *RAIN*, we used the rain function with the parameters “period=17”, “period.delta=13”, and *RAIN*’s adaptive Benjamini-Hochberg multiple testing correction. Cycling genes had an adjusted p-value<0.05. Functional enrichment analysis of cyclic genes was performed with R package *gprofiler2* v0.2.3, with the g:SCS multiple testing correction method^63, 64^. The databases queried with *gprofiler2* included Gene Ontology, KEGG, Reactome, WikiPathways, TRANSFAC, miRTarBase, the Human Protein Atlas, CORUM, and Human Phenotype Ontology. Finally, we found the overlap between *MetaCycle* rhythmic genes and proximal/eQTL-mapped genes corresponding to 236 putative causal single nucleotide polymorphisms that Jones *et al.* previously associated with self-reported chronotype in humans^24^.

### Integrating mRNA and miRNA pairwise

To discover relationships between cycling mRNAs and miRNAs, we used R packages *lmms* v1.3.3 and *dynOmics* v1.0 to collapse replicates and find the delayed Pearson correlations between rhythmic mRNA-miRNA pairs^65, 66^. After accounting for the predicted delay in log2 expression, significant associations had a Benjamini-Hochberg adjusted p-value<0.05. We considered associations with a negative correlation and time delay≤0, i.e., decreased mRNA expression after miRNA expression, to be potential miRNA direct targets. These associations were compared to predicted and/or experimentally confirmed mRNA targets of cycling miRNAs identified with R package *multiMiR* v1.26.0^67^. Before querying these databases, the precursor miRNA IDs were converted to their respective mature versions with R package *miRBaseConverter* v1.11.1^68^.

### Identifying co-expression networks of mRNA and miRNA

We used weighted gene co-expression network analysis (R package *WGCNA* v1.72.5) to find interactions between groups of co-expressed mRNA and miRNAs^69^. We calculated the pairwise Pearson correlations between the log2 transformed gene counts for each tissue. We calculated gene similarities by adding 1 to each correlation and dividing by 2, thus creating a signed network. Using various soft-thresholding powers, we created weighted adjacency matrixes with scale-free topology fits (R^2^) > 0.8. mRNA powers: 16 (cortex), 16 (hypothalamus), 18 (striatum), 16 (liver). miRNA powers: 4 (cortex), 5 (hypothalamus), 6 (striatum), and 9 (liver). We then calculated topological overlap matrixes (TOM). Based on dissimilarity (1-TOM), genes were grouped into modules by hierarchical clustering with the dynamic tree-cut algorithm.

Once the networks were constructed, we found each module’s hub gene, which is the gene with the highest connectivity within a module. We also used a hypergeometric test to determine the enrichment of cycling genes, both from *MetaCycle* and *RAIN*. The background set for all hypergeometric tests was the genes remaining after non-specific filtering. We then used the Spearman correlation between mRNA and miRNA “cycling” modules’ eigengenes to find potential mRNA-miRNA relationships, with p-values adjusted using the Benjamin-Hochberg method. We also used a hypergeometric test to determine the enrichment of brain cell types in cycling modules. Yao *et al.*’s single-cell sequencing data from the mouse cortex and hippocampus provided the gene-expression profiles of cell types^70^. For each cell type subclass, we considered a marker gene identified by Yao *et al.* to be expressed if its trimmed mean expression is greater than zero. We then found the overlap between genes expressed in a cell type subclass and a module. The grey60 module in the cortex was visualized with *Cytoscape* 3.10.0. Finally, we performed pathway analysis of modules with R package *gprofiler2,* as described above.

### Animals for immunohistochemistry

All work was performed using 20-to 30-week-old C57BL/6J male mice, matching the animals’ age in the miRNA/mRNA-seq datasets. The institutional animal care and use committees of Queen’s University approved all animal care and procedures (Protocol # 2023–2428) following all guidelines of the Canadian Council on Animal Care.

### Immunohistochemistry

Animals were transcardially perfused with phosphate buffered saline (PBS), followed by 4% paraformaldehyde (PFA). The brain was removed and placed into 30% sucrose in PBS for 24 h, embedded in Tissue TEK OCT (Fisher Scientific) and frozen at -80°C. 16 μm sections were collected using a Leica CM1950 cryostat (Leica). Sections were blocked with 4% BSA (Sigma-Aldrich) and 0.3% Triton X-100 (BioShop) in PBS for 1h at room temperature and incubated overnight at 4°C with primary antibodies. Primary antibodies for proteins of interest were anti-IL-33 (1:50, cat.no.AF3626, R&D Systems), anti-AdipoR2 (1:200, cat.no. BS-0611R, Bioss), anti-FZD4 (1:200, cat.no. SAB4503265, Sigma-Aldrich), anti-EIF1B(1:100, cat.no. ABX326521, Abbexa Ltd), anti-LRRK2 (1:200,cat.no.ab133474, Abcam), anti-SPHK2(1:200, cat.no.17096-1-AP, Proteintech), and anti-Transferrin (1:50, cat.no.ZRB1225, Sigma-Aldrich). For markers of neurons, microglia, astrocytes, and oligodendrocytes, anti-NeuN (1:200, cat.no MAB377, Sigma-Aldrich), anti-Iba1 (1:500, cat.no. 019-19741, Wako; 1:500 cat.no. 234 009, Synaptic Systems), anti-GFAP (1:1000, cat.no.13-0300, Invitrogen;1:1000), anti-Olig2 (1:250, cat.no. MABN50, Sigma Aldrich). After washing, sections were incubated with either Alexa Fluor 488, 594, or 647 conjugated secondary antibodies (1:500, Invitrogen) overnight at 4°C. The slides were mounted using Prolong™ Diamond Antifade Mountant with DAPI (Life Technologies).

### Image Analysis

Images were captured using an Eclipse Ti2 microscope (Nikon) with NIS-Elements AR (Nikon). Five x40 images were captured in the brain region of interest for each animal, and for each protein and CNS cell marker staining combination. ImageJ (NIH) was used to quantify the fluorescence intensity of each protein within the boundaries of each CNS cell body. Multichannel images were split into single channels, and then channels containing CNS cell markers were thresholded to only contain fluorescence signals within the cell body. The thresholded image was turned into a selection, which created a region of interest (ROI) around the cells. The channel containing the hub gene protein was converted to a 16-bit image so that pixel ranges were consistent, and the mean gray value of the image was measured. Then, the ROI’s were superimposed onto the channel, and the mean gray value of the ROI was measured. A circular ROI with a diameter of 20µm (area 316.204µm^2^) was created and placed in a region of the image without hub gene protein signal to measure background fluorescence. To calculate fluorescence intensity, the mean background fluorescence was subtracted from the mean fluorescence intensity from the CNS cell ROI. The fluorescence intensity for each image was averaged together.

### Statistics and Reproducibility

Unless otherwise described, transcriptomics data analyses were conducted in R v4.4.0. Immunochemistry analyses used Graphpad Prism (v10, Graphpad Software, Boston, USA). Analysis of variance (ANOVA) was used to compare protein fluorescence data across timepoints and cell types, with significance thresholds set at *P*<0.05, using the post hoc Tukey test and Holm-Sídak multiple comparisons test.

## Data availability

Bulk sequencing data are available at <u>GSE151567</u>. Cycling genes and mRNA-miRNA pairs can be interactively explored at https://www.ghasemloulab.ca/chronoCNS. All other data supporting this study’s findings are available from the corresponding authors upon reasonable request.

## Code availability

No custom code was used to generate or process the data. However, upon publication, scripts used to analyze data and generate figures will be available at https://github.com/ComputationalGenomicsLaboratory/chronoCNS.

## Extended Data

**Extended Data Fig. 1:**
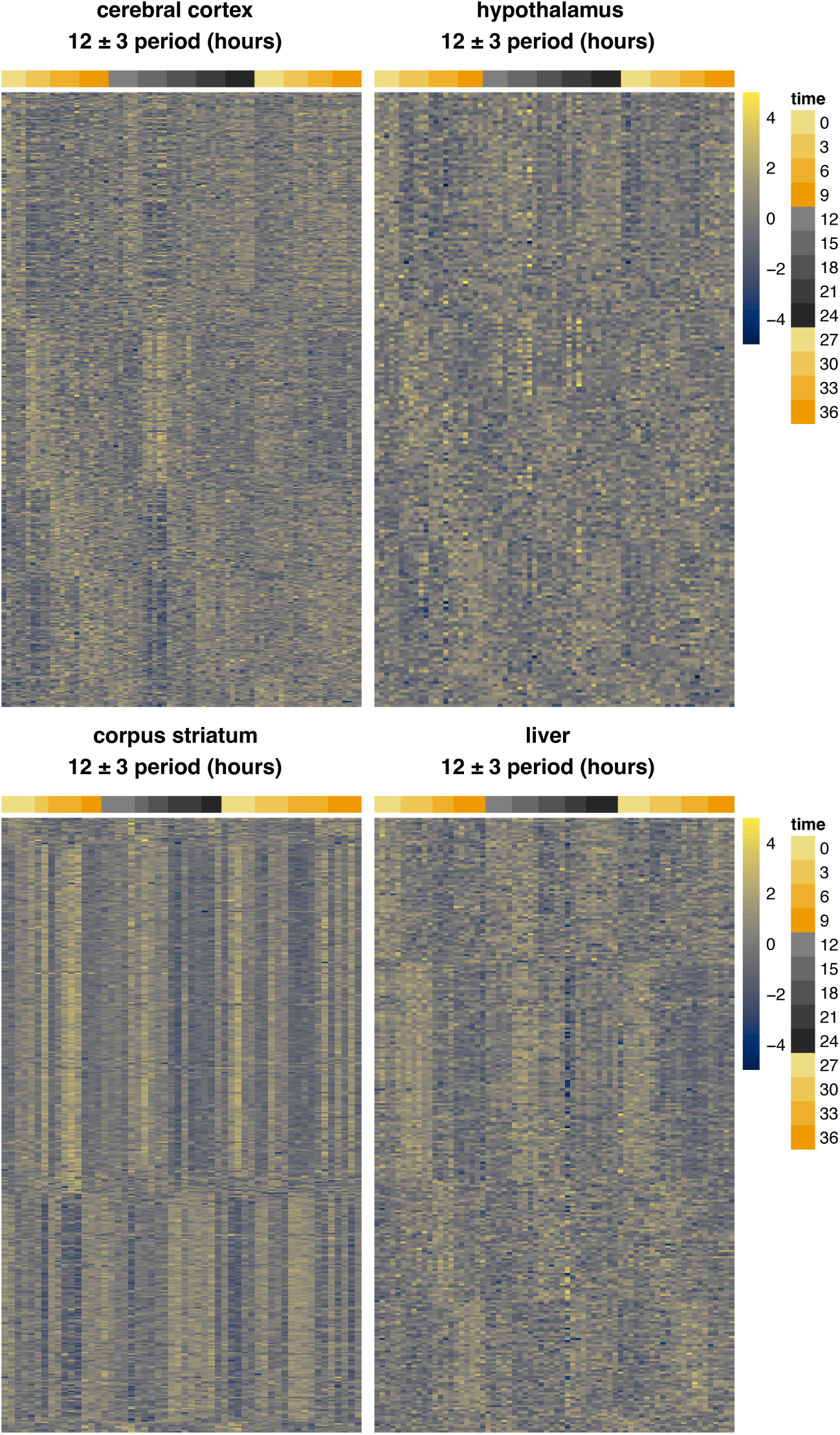
Expression of mRNA cycling genes with a period length in hours 12±3. Heatmaps are labelled by their respective tissues. Rows are genes, and columns are samples. Genes are sorted by phase. Samples are annotated by their time of collection, measured in Zeitgeber Time. Colours represent gene counts transformed by the *DESeq2* variance stabilizing transformation.

**Extended Data Fig. 2:**
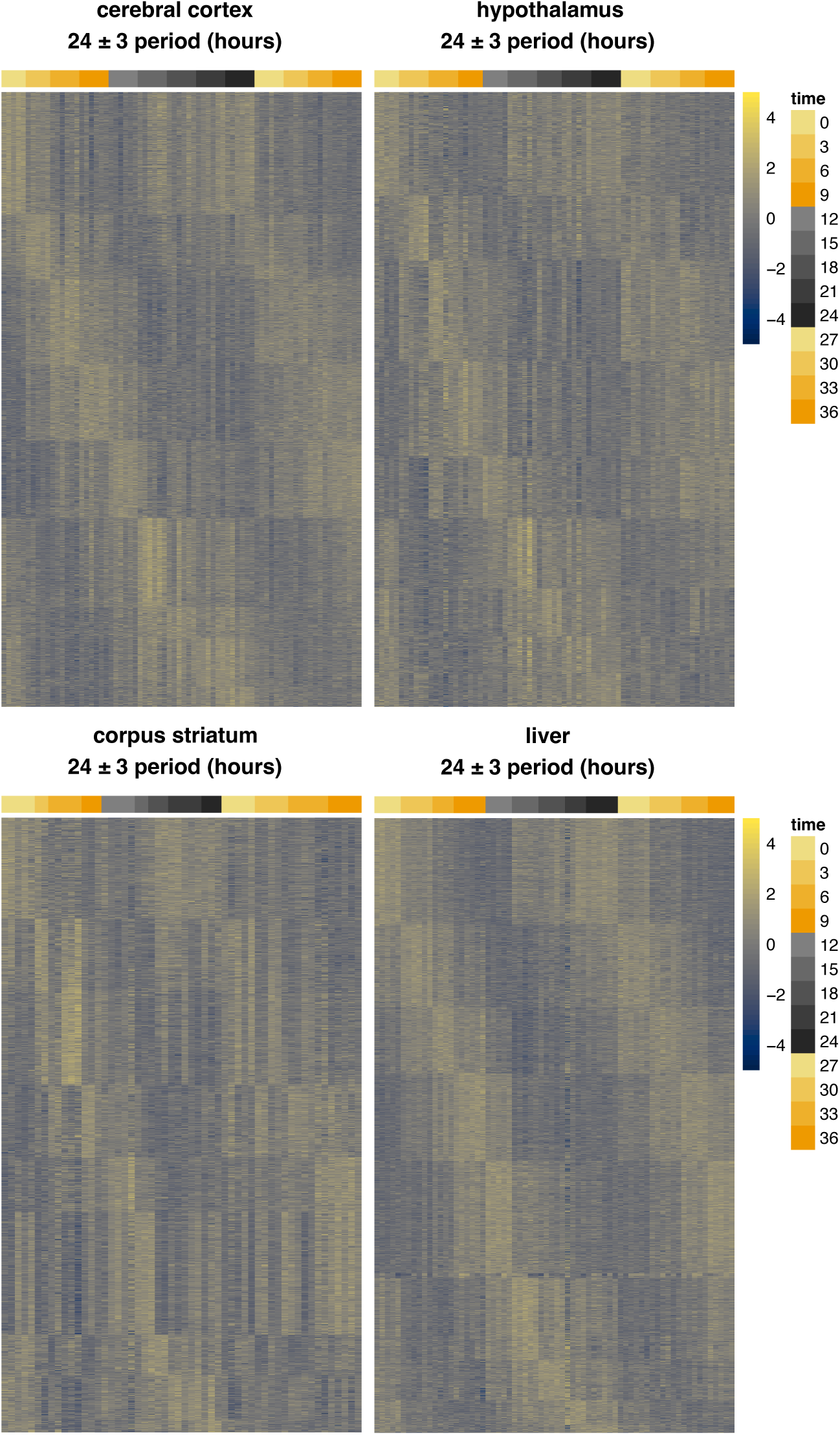
Expression of mRNA cycling genes with a period length in hours 24±3. Heatmaps are labelled by their respective tissues. Rows are genes, and columns are samples. Genes are sorted by phase. Samples are annotated by their time of collection, measured in Zeiteber Time. Colours represent gene counts transformed by the *DESeq2* variance stabilizing transformation.

**Extended Data Fig. 3:**
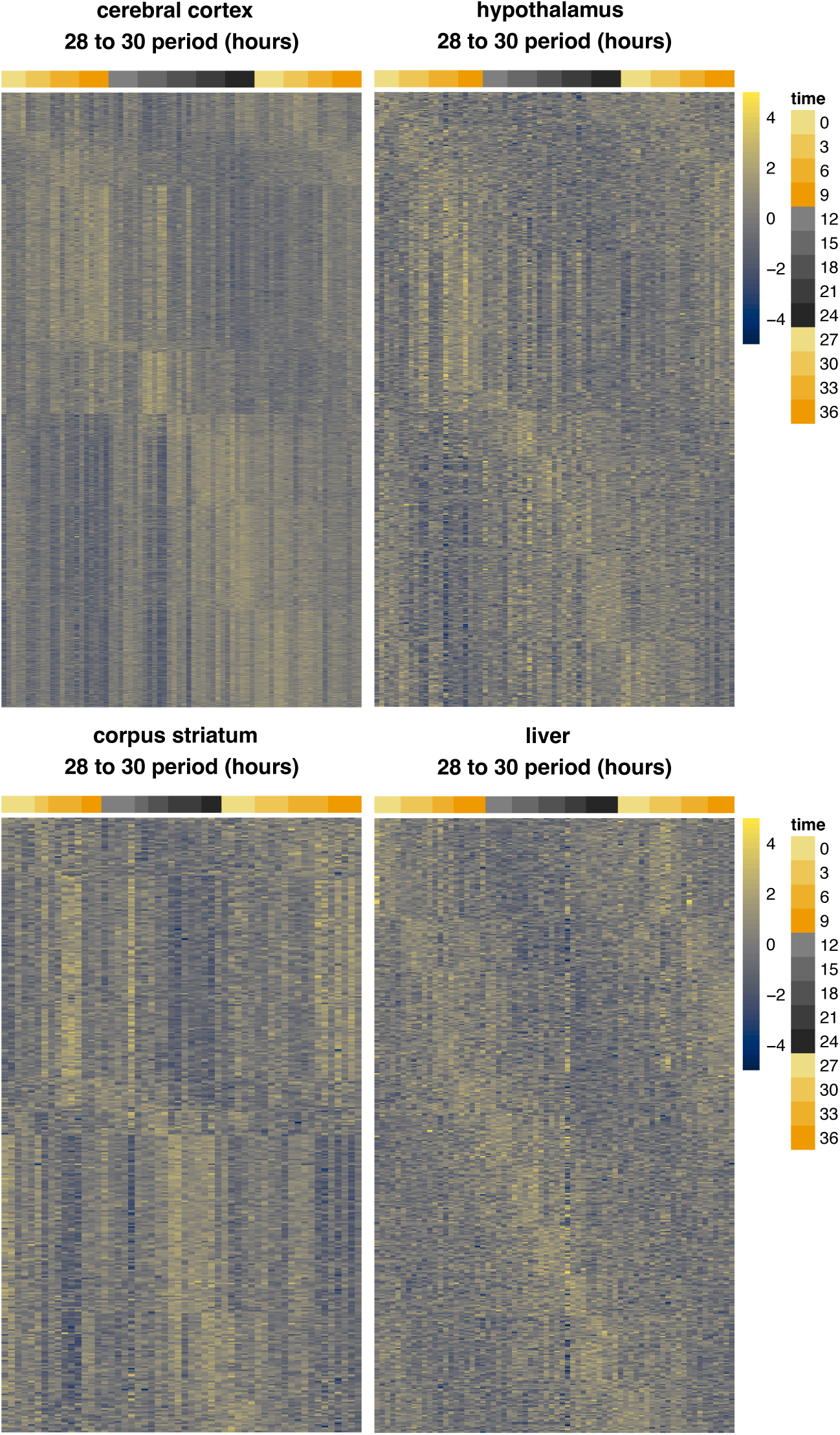
Expression of mRNA cycling genes with a period length in hours >27 and ≤30. Heatmaps are labelled by their respective tissues. Rows are genes, and columns are samples. Genes are sorted by phase. Samples are annotated by their time of collection, measured in time. Colours represent gene counts transformed by the *DESeq2* variance stabilizing transformation.

**Extended Data Fig. 4:**
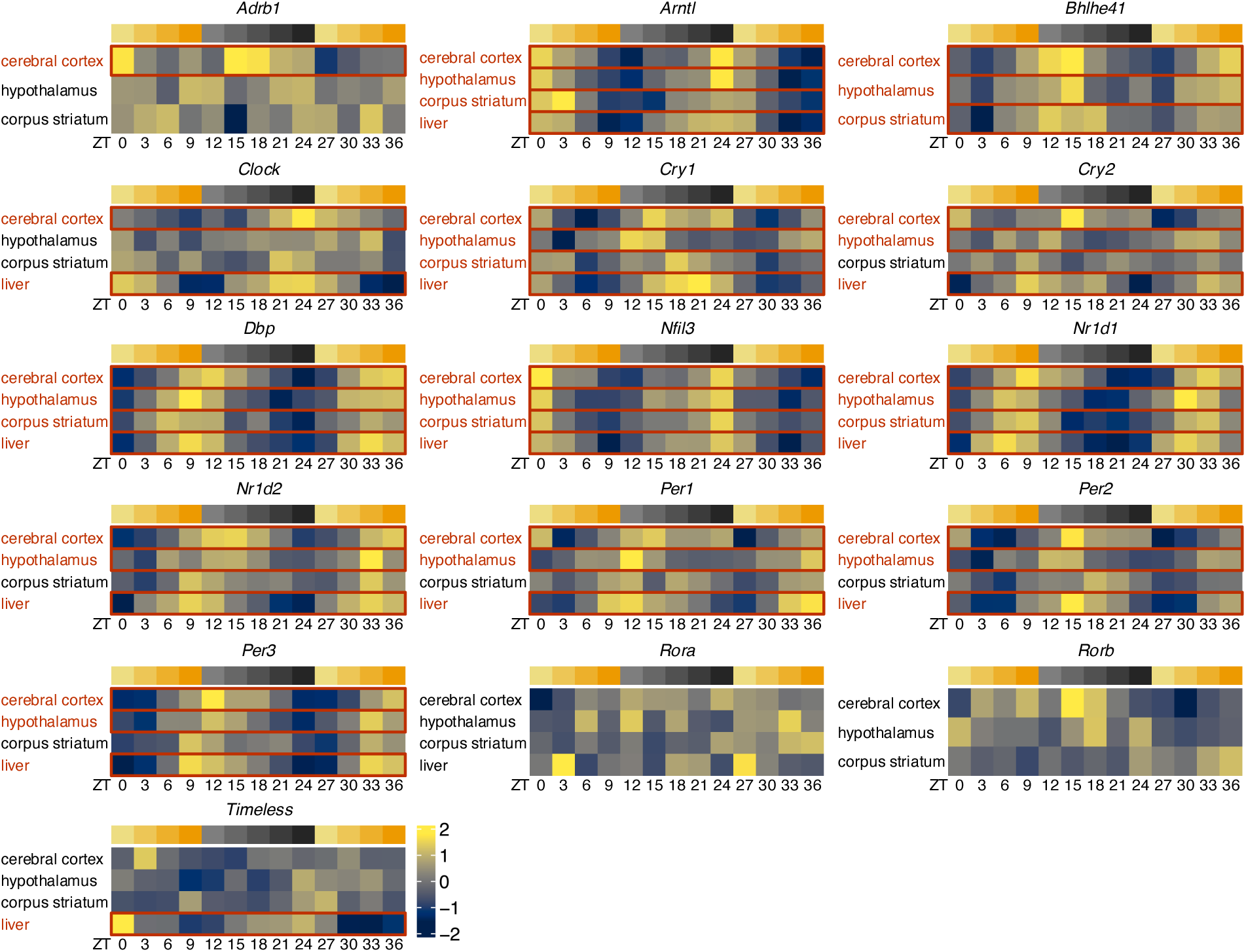
Expression of key clock genes across time and sites. Heatmaps are labelled by their respective genes. Rows are tissues, and columns are timepoints measured in Zeitgeber Time (ZT). Red text and boxes indicate that a clock gene is cycling in its respective tissue (*MetaCycle* P_BH_<0.05). Variance stabilizing transformed gene counts are center and standard deviation scaled by gene. These scaled counts are then averaged across samples from the same timepoint and tissue and are represented by colours ranging from yellow to blue.

**Extended Data Fig. 5:**
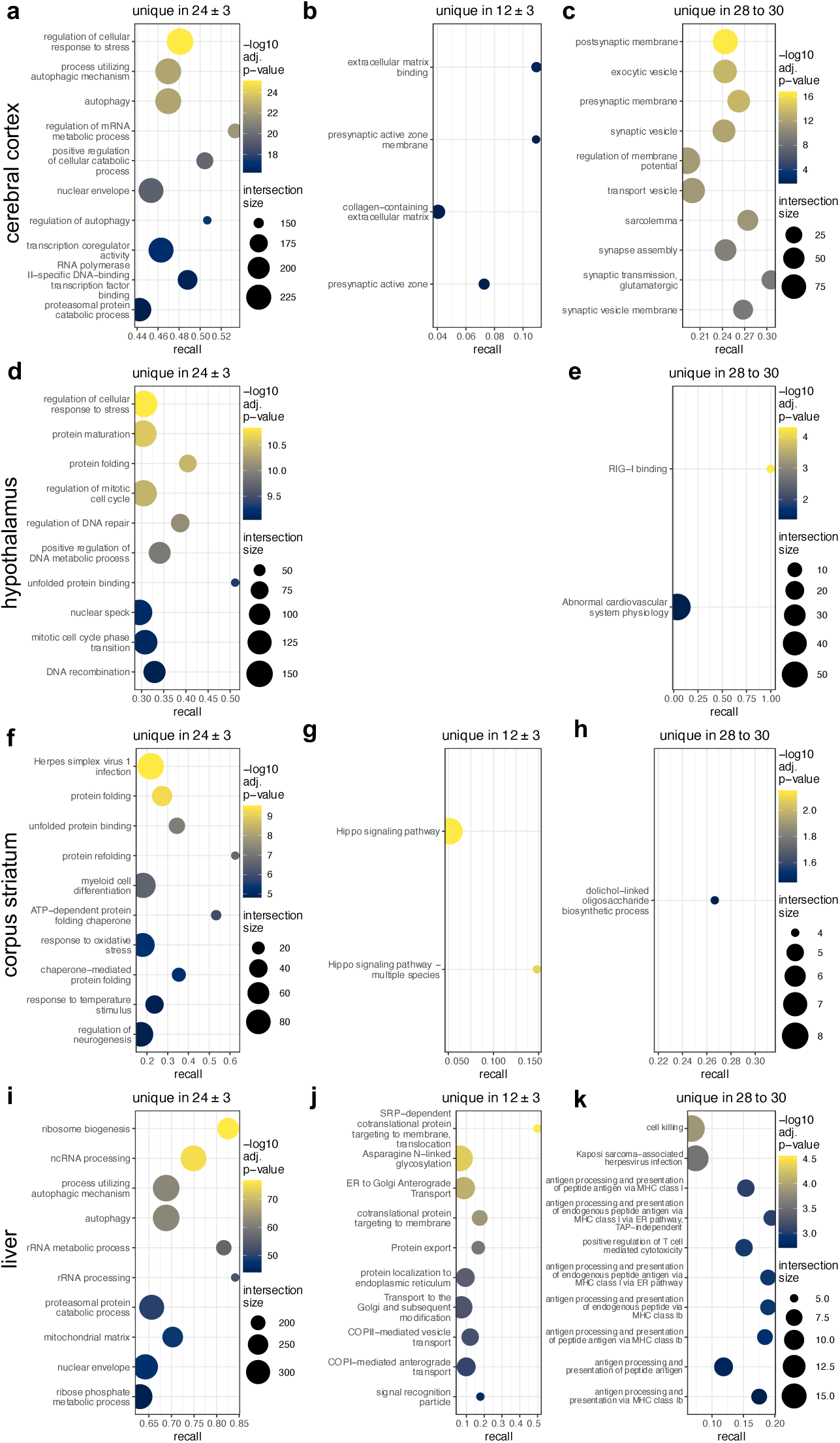
Comparison of pathways enriched in cycling genes across period length categories (24-hour, 12-hour, and 28-to 30-hours). Top pathways that are uniquely enriched in the 3 period categories for each tissue (*P*_g:SCS_<0.05; 10≤term size ≤500). Intersection size is the number of genes that are both cycling and in a pathway. Recall is intersection size divided by the size of a term/pathway. a-c, Comparison of pathways within the cerebral cortex. d,e, Comparison of pathways within the hypothalamus. No pathways were unique to the period category 12 ± 3. For e, results are not filtered by term size. f-h, Comparison of pathways within the corpus striatum. i-k, Comparison of pathways within the liver.

**Extended Data Fig. 6:**
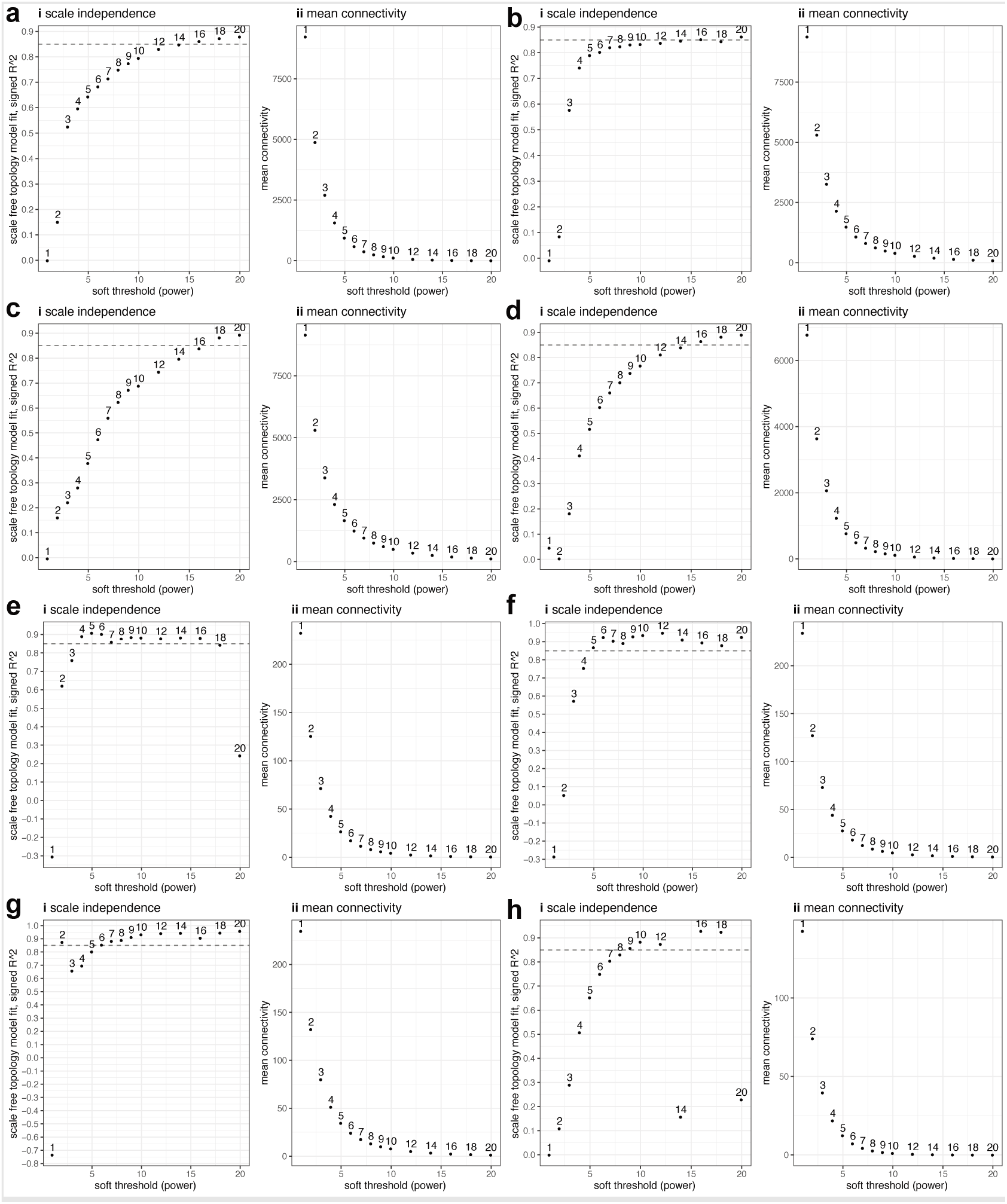
Soft thresholding plots used for network construction. X-axes are the soft thresholding powers considered. **a-d,** Plots for mRNA datasets. Plots are in the order of 1) cortex, 2) hypothalamus, 3) striatum, and 4) liver. **e-h,** Plots for microRNA datasets. Plots are in the order of 1) cortex, 2) hypothalamus, 3) striatum, and 4) liver. Within each panel, there are i) a scale independence plot and ii) a mean connectivity plot. For **i,** the y-axis is the scale-free topology model fit, which measures how similar the network’s topology is to an ideal scale-free topology. For **ii,** the y-axis is the average of how correlated each gene in the network is to all other network genes.

**Extended Data Fig. 7:**
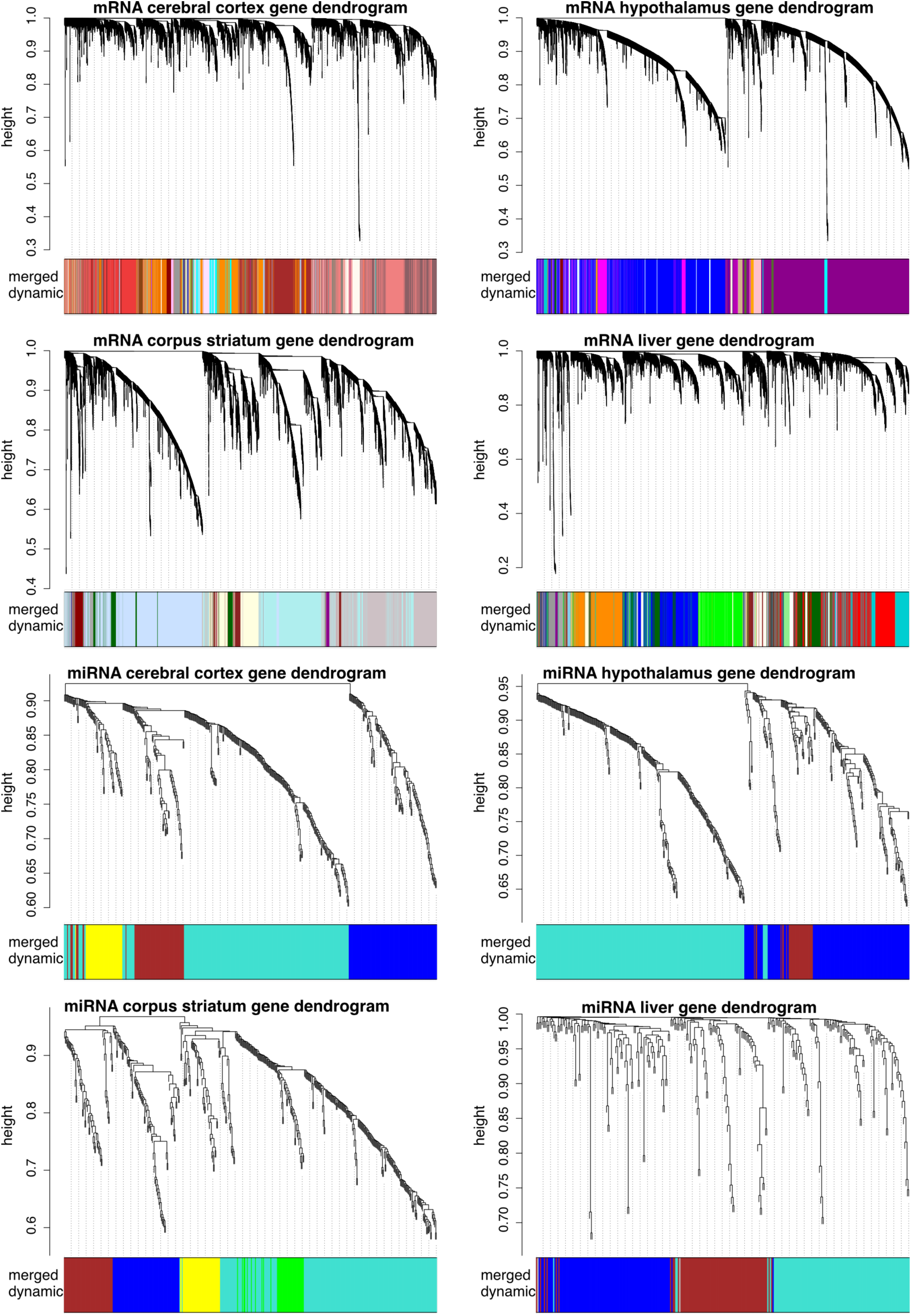
Gene dendrograms representing the networks constructed for each sequencing type and tissue. Dendrograms are labelled by their respective sequencing types (mRNA or miRNA) and tissues. Modules of genes are indicated by colours displayed at the bottom of each dendrogram.

**Extended Data Fig. 8:**
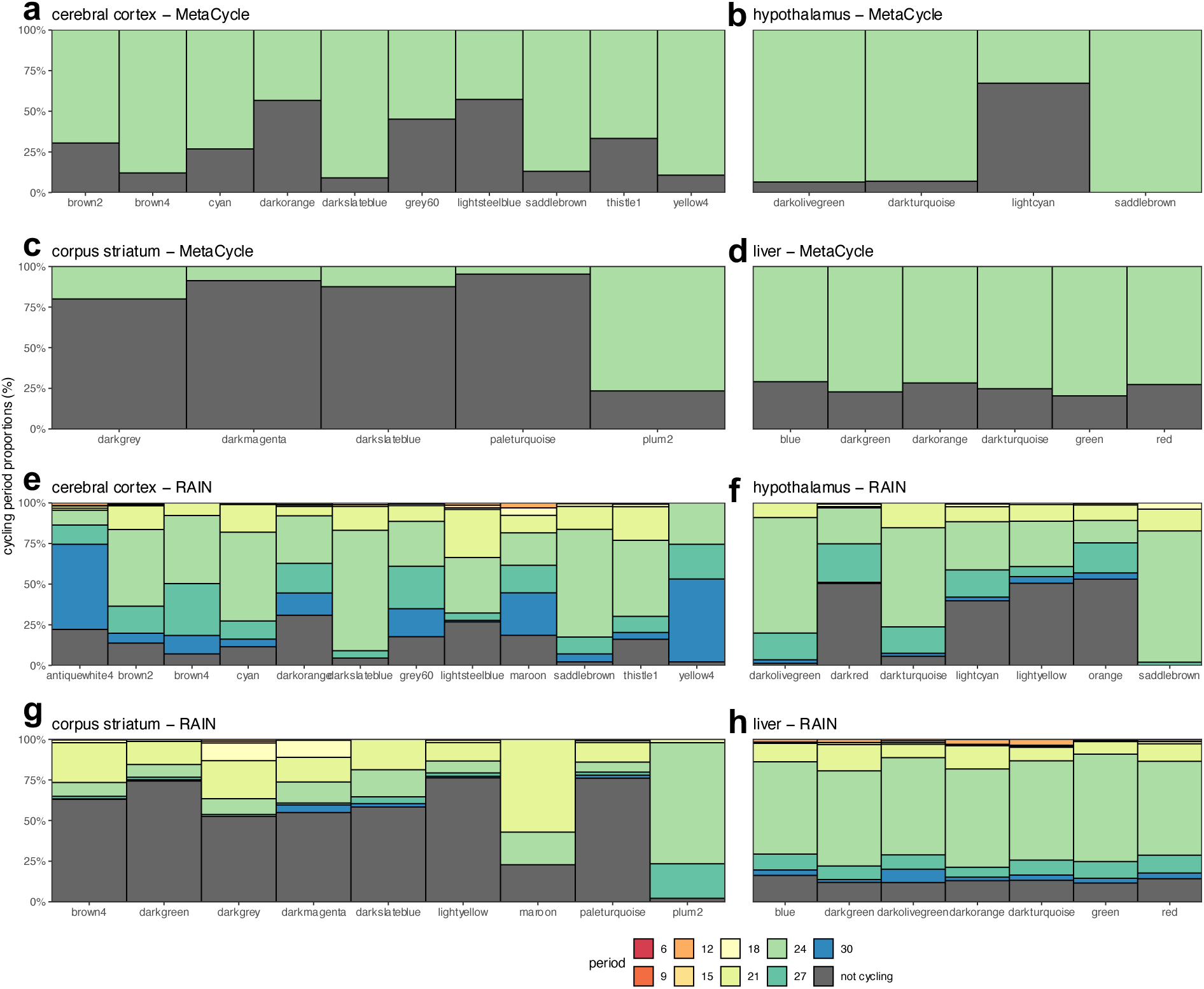
Composition of mRNA periods within cycling mRNA modules. Modules are on the x-axes. The y-axes indicate the number of cycling genes with a certain period in a module divided by the total number of genes in a module and multiplied by 100. **a-d,** Modules enriched in cycling mRNAs identified by *MetaCycle*. Periods are estimated by *MetaCycle*. **e-g,** Modules enriched in cycling mRNAs identified by *RAIN*. Periods are estimated by *RAIN*.

**Extended Data Fig. 9:**
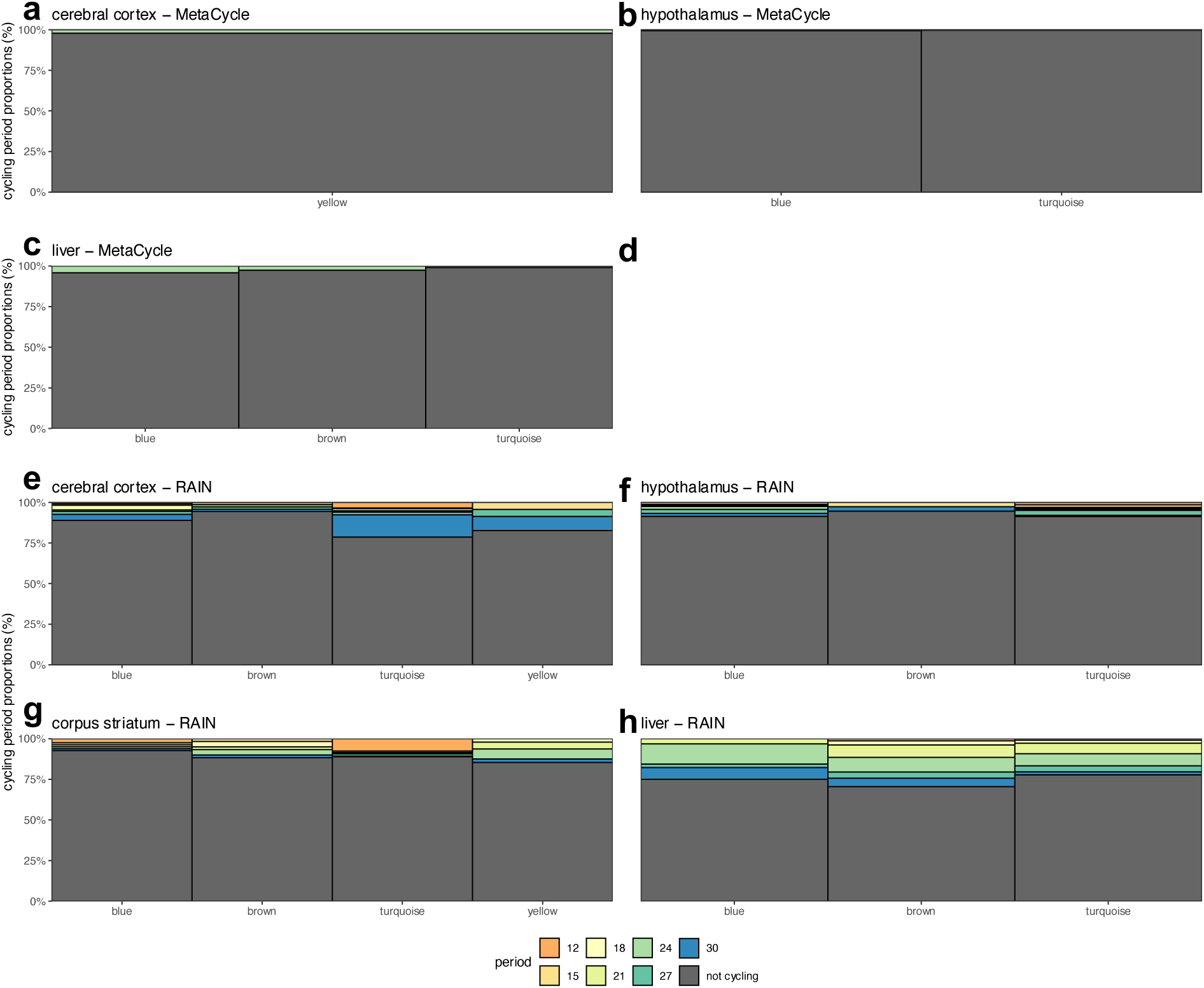
Composition of miRNA periods within cycling miRNA modules. Modules are on the x-axes. The y-axes indicate the number of cycling genes with a certain period in a module divided by the total number of genes in a module and multiplied by 100. **a-d,** Modules containing cycling miRNAs identified by *MetaCycle*. Periods are estimated by *MetaCycle*. **e-g,** Modules containing in cycling miRNAs identified by *RAIN*. Periods are estimated by *RAIN*. In the text, there are Spearman correlation coefficients, and Benjamini-Hochberg adjusted P-values in parentheses. miRNA modules are on the x-axes, and mRNA modules are on the y-axes. Cells in heatmaps are filled by the Spearman correlation coefficients (ρ).

**Extended Data Fig 10.**
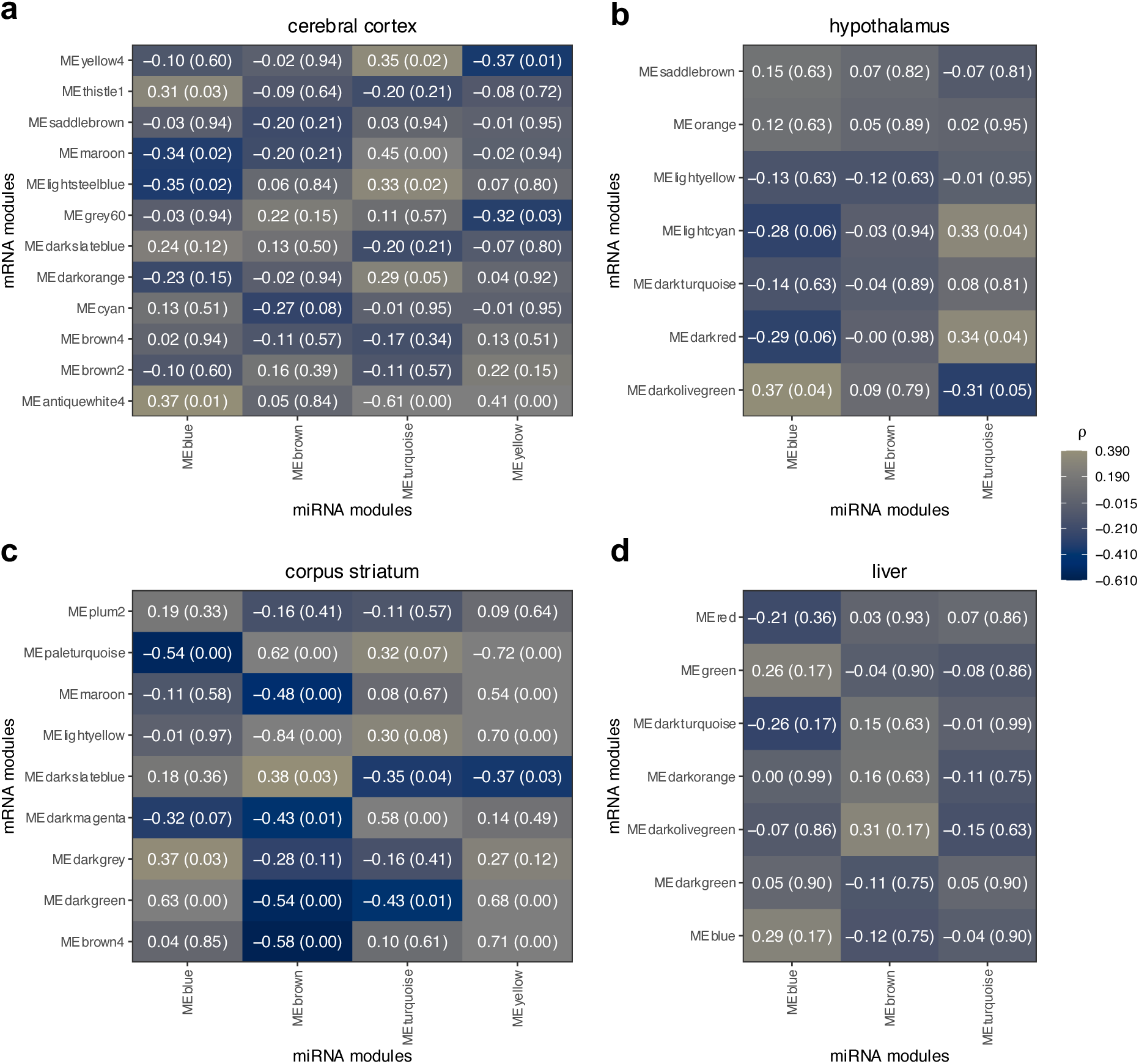
Correlation between cycling mRNA and miRNA module eigengenes. In the text, there are Spearman correlation coefficients, and Benjamini-Hochberg adjusted P648 values in parentheses. miRNA modules are on the x-axes, and mRNA modules are on the y-axes. Cells in heatmaps are filled by the Spearman correlation coefficients (ρ).

**Extended Data Fig. 11:**
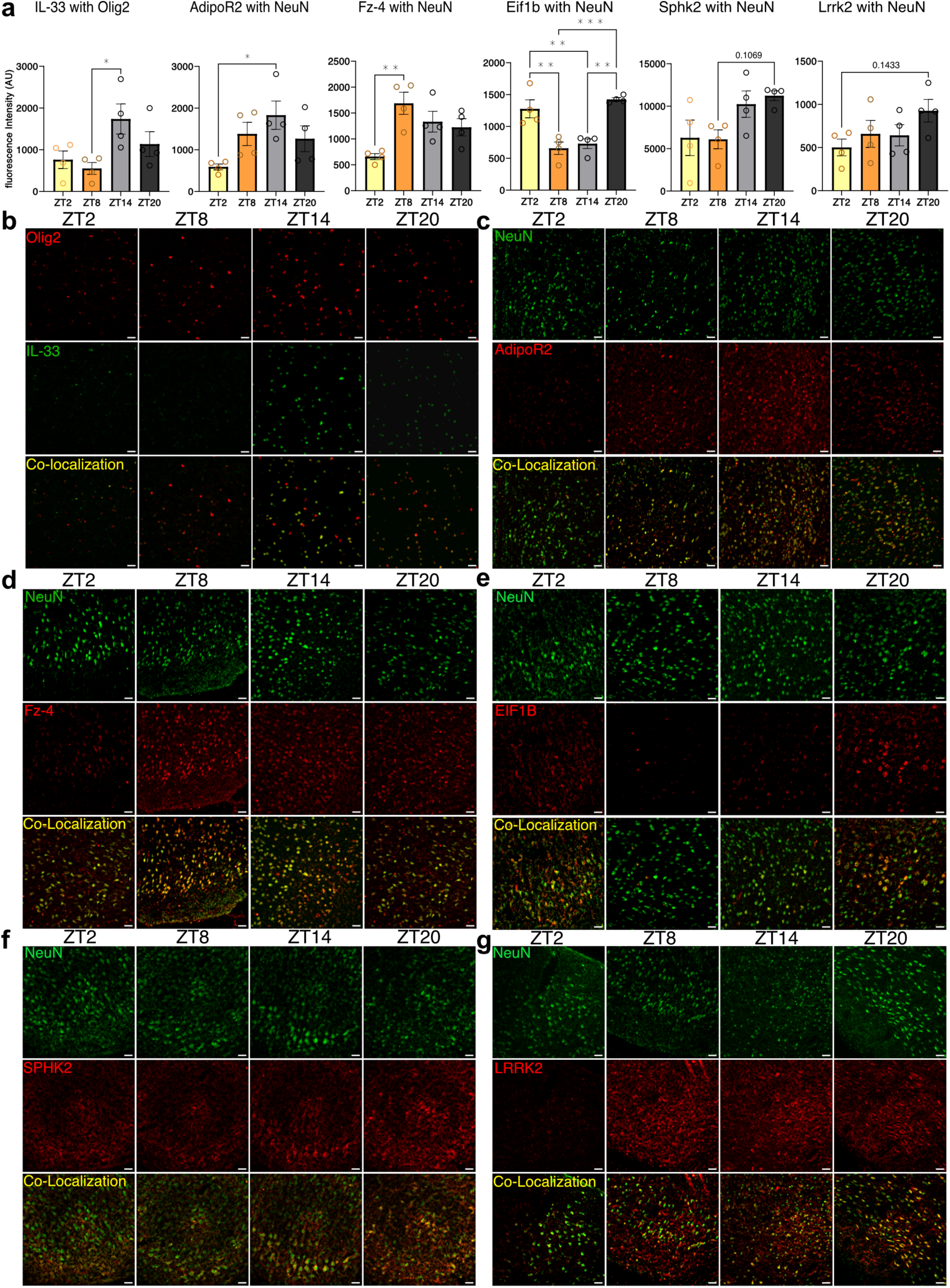
Expression of candidate proteins by neurons in the cortex and hypothalamus. **a**, Quantification of fluorescence intensity of IL-33 colocalized with oligodendrocyte marker Olig2 in the cortex; AdipoR2 co-localized with neuronal marker NeuN in the hypothalamus; Fz-4, Sphk2, Eif1b, and Lrrk2 co-localized with NeuN in the cortex, in brains collected at ZT2, ZT8, ZT14, and ZT20. One-way ANOVA with Tukey’s post hoc test, *n*=4 animals for each timepoint. IL-33 with Olig2: ZT8 versus ZT14, \**P*=0.0371, *q*=4.443, d.f.=12. AdipoR2 with NeuN: ZT2 versus ZT14, \**P*=0.0313, *q*=4.579, d.f.=12; Fz-4 with NeuN: ZT2 versus ZT8, \*\**P*=0.0053, *q*=6.027, d.f.=12; Eif1b with NeuN: ZT8 versus ZT20, \*\*\**P*=0.0005, *q*=8.092, d.f.=12, ZT2 versus ZT8, \*\**P*=0.0029, *q*=6.521, d.f.=12, ZT2 versus ZT14, \*\**P*=0.0072, *q*=5.769, d.f.=12, ZT14 versus ZT20, \*\**P*=0.0011, *q*=7.340, d.f.=12; Sphk2 with Neun: ZT8 versus ZT20, *P*=0.1069, *q*=3.563, d.f.=12; Lrrk2 with Neun: ZT2 versus ZT20, *P*=0.1433, *q*=3.308, d.f.=12. All data presented as mean±s.e.m. **b,** Representative images of hypothalamic neurons (labelled with NeuN), AdipoR2, and their co-localization at ZT2, ZT8, ZT14, and ZT20. Scale bar, 25µm. **c,** Representative images of cortical neurons, Fz-4, and their co-localization at ZT2, ZT8, ZT14, and ZT20. Scale bar, 25µm. **d,** Representative images of cortical neurons, Eif1b, and their co-localization at ZT2, ZT8, ZT14, and ZT20. Scale bar, 25µm. **e,** Representative images of cortical neurons, Sphk2, and their co-localization at ZT2, ZT8, ZT14, and ZT20. Scale bar, 25µm. **f,** Representative images of cortical neurons, Lrrk2, and their co-localization at ZT2, ZT8, ZT14, and ZT20. Scale bar, 25µm.

**Extended Data Fig. 12:**
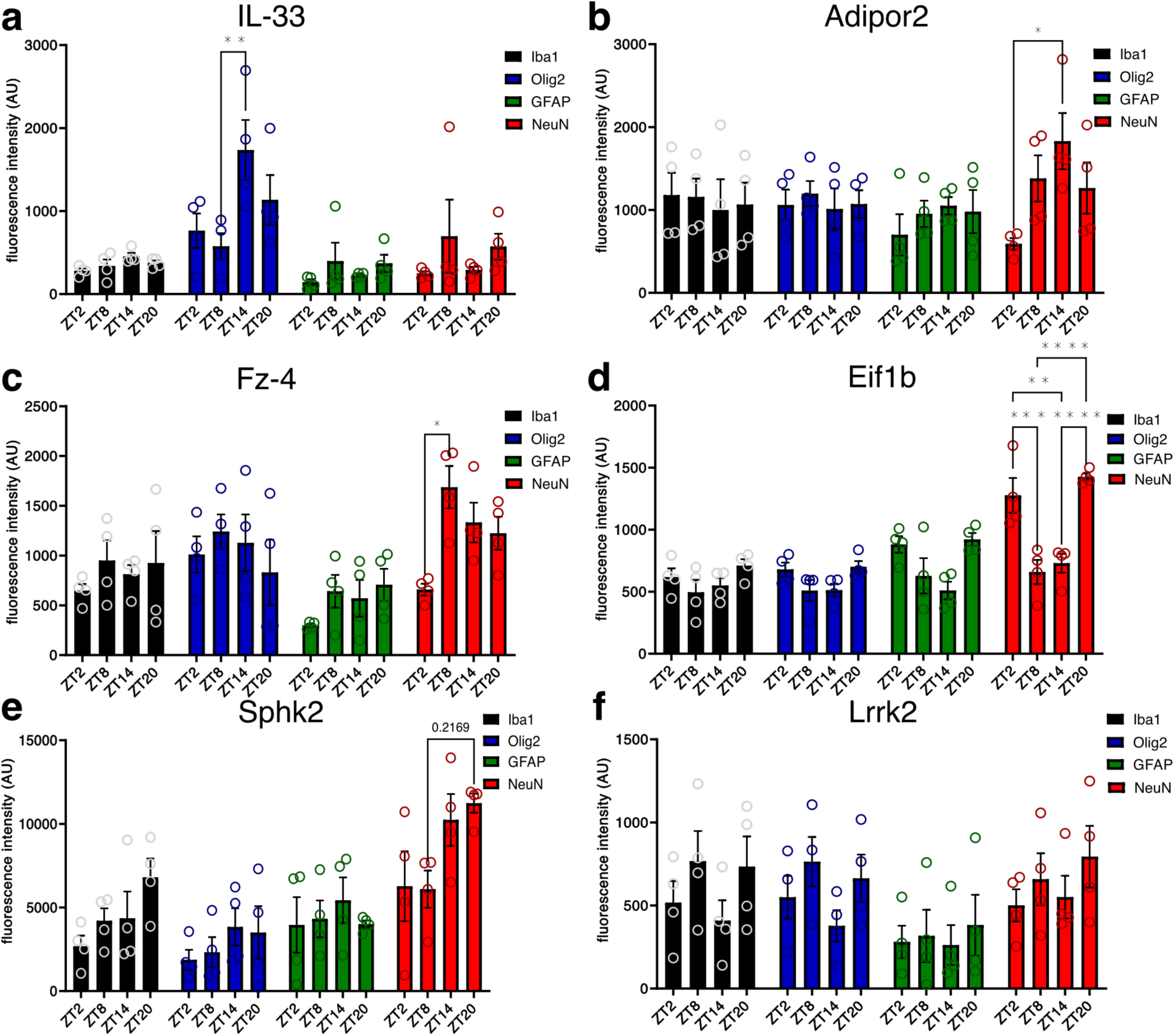
Quantified expression of candidate proteins in microglia, oligodendrocytes, astrocytes, and neurons. **a-f**, Quantification of fluorescence intensity of candidate proteins colocalized with Iba1 (microglia), Olig2 (oligodendrocyte marker), GFAP (astrocytes), and NeuN (neurons) at ZT2, ZT8, ZT14, and ZT20. Two-way ANOVA, with Tukey’s post hoc test, *n*=4 animals for each timepoint. All data presented as mean ± s.e.m. **a,** IL-33 with Olig2 in cortex, ZT8 versus ZT14, \*\**P*=0.0484, *q*=5.127, d.f.=48. **b,** AdipoR2 with NeuN in hypothalamus: ZT2 versus ZT14, \**P*=0.0313, *q*=4.579, d.f.=12. **c,** Fz-4 with NeuN in cortex: ZT2 versus ZT8, \**P*=0.0379, *q*=5.259, d.f.=48, **d,** Eif1b with NeuN in cortex: ZT2 versus ZT8, \*\*\**P*=0.0002, *q*=7.622 d.f.=48, ZT2 versus ZT14 \*\**P*=0.0017, *q*=6.744, d.f.=48, ZT8 versus ZT20 \*\*\*\**P*<0.0001, *q*=9.458, d.f.=48, ZT14 versus ZT20 \*\*\*\**P*<0.0001, *q*=8.579, d.f.=48. **e,** Sphk2 with NeuN in cortex: ZT8 versus ZT20, *P*=0.2169, *q*=4.211, d.f.=48. **f,** Lrrk2 with NeuN in cortex: ZT2 versus ZT20, *P*=0.9846, *q*=2.031, d.f.=48.

## Acknowledgements

This work was supported by grants from the Natural Sciences and Engineering Research Council of Canada (NSERC; #RGPIN-05604, to N.G.) and Brain Canada (to N.G.). Computations were performed on resources and with support provided by the Centre for Advanced Computing (CAC) at Queen’s University in Kingston, Ontario. The CAC is supported in part by funding from Queen’s University, the Digital Research Alliance of Canada and the Government of Ontario. The funders had no role in study design, data collection and analysis, decision to publish, or preparation of the manuscript.

## Author contributions

A.Z., C.O., D.G.T., H.C., H.G., Q.L.D., and N.G. conceived and planned analyses. Computational analyses were performed by A.Z, with support from Z.Y.F. Experiments were performed and analyzed by C.O. Interpretation of findings was conducted by A.Z., C.O., Q.L.D., and N.G. Paper preparation was largely performed by A.Z., C.O., and N.G. The figures were prepared by A.Z. and C.O. All authors reviewed the results and contributed to the final paper.

